# Gene set enrichment analysis for genome-wide DNA methylation data

**DOI:** 10.1101/2020.08.24.265702

**Authors:** Jovana Maksimovic, Alicia Oshlack, Belinda Phipson

**Affiliations:** Peter MacCallum Cancer Centre, Melbourne, Victoria, 3000, Australia; Department of Pediatrics, University of Melbourne, Parkville, Victoria, 3010, Australia; Murdoch Children’s Research Institute, Parkville, Victoria, 3052, Australia; School of Biosciences, University of Melbourne, Parkville, Victoria, 3010, Australia; Sir Peter MacCallum Department of Oncology, University of Melbourne, Parkville, Victoria, 3010, Australia

**Keywords:** DNA methylation, gene set analysis, differential methylation, statistical analysis

## Abstract

DNA methylation is one of the most commonly studied epigenetic marks, due to its role in disease and development. Illumina methylation arrays have been extensively used to measure methylation across the human genome. Methylation array analysis has primarily focused on preprocessing, normalisation and identification of differentially methylated CpGs and regions. GOmeth and GOregion are new methods for performing unbiased gene set testing following differential methylation analysis. Benchmarking analyses demonstrate GOmeth outperforms other approaches and GOregion is the first method for gene set testing of differentially methylated regions. Both methods are publicly available in the *missMethyl* Bioconductor R package.

## Background

DNA methylation is essential to human development, with roughly 3-6% of all cytosines methylated in normal human DNA (Esteller 2007). Epigenetic marks can be modified by environmental exposures, and methylation changes are known to accumulate with age. Aberrant methylation patterning is associated with many diseases, which has led to several large studies profiling DNA methylation, such as The Cancer Genome Atlas, Encyclopedia of DNA Elements and numerous epigenome-wide association studies.

Both array and sequencing based technologies are available for profiling DNA methylation at a genome-wide scale. However, even though the cost of sequencing has dramatically decreased, the ease and cost effectiveness of the Illumina human methylation arrays have ensured that the array platforms remain a popular choice. To date, a major focus when analysing DNA methylation data has been the identification of significantly differentially methylated CpG sites between groups of samples in a designed experiment. There are many well-established analysis methods that perform normalisation and statistical testing for this purpose, including publicly available software packages, such as *limma* (Ritchie et al. 2015), *minfi* (Aryee et al. 2014), *missMethyl* (Phipson, Maksimovic, and Oshlack 2016), *methylumi* (Davis et al. 2019), *wateRmelon* (Pidsley et al. 2013), *ChAMP* (Morris et al. 2014), *RnBeads* (Assenov et al. 2014; Müller et al. 2019), *Harman* (Oytam et al. 2016), and *ENmix* (Z. Xu et al. 2016). It is well established that methylation of CpG sites is spatially correlated along the genome (Eckhardt et al. 2006) and that long tracks of differential methylation are often more biologically meaningful than differences at individual CpG sites (Hansen et al. 2011). This has led to region-based analyses, with Bioconductor R packages such as *Probe lasso* (Butcher and Beck 2015), *bumphunter* (Jaffe et al. 2012), *DMRcate* (Peters et al. 2015), *mCSEA* (Martorell-Marugán, González-Rumayor, and Carmona-Sáez 2019) and *DMRforPairs* (Rijlaarsdam et al. 2014) developed specifically for this purpose.

Once differential methylation analysis between groups of samples has been performed, there may be a long list of significant CpG sites or regions for the researcher to interpret. A popular approach to gain a more systems-level understanding of the changes in methylation is to examine which gene pathways may be enriched for differential methylation in the experiment. This approach was popularised in the analysis of gene expression microarrays and RNA-sequencing (RNA-Seq) data, with one of the first methods, GSEA, published in 2005 (Subramanian et al. 2005). Since then, a number of gene set testing methods have been developed (e.g. Yaari et al. (2013), Wu et al. (2010), and Wu and Smyth et al. (2012)), all of which are specific to gene expression data, with the GOSeq method (Young et al. 2010) developed specifically to account for gene length bias in RNA-seq data.

Methylation, however, is a DNA mark that can occur anywhere on the genome and is not as directly related to genes as expression data. Therefore, a methylation specific issue in performing gene set testing is how to assign differentially methylated features to genes. Thus far, there are very few gene set testing methods designed specifically for DNA methylation data, and often ad hoc approaches are taken. Only two other methods, ebGSEA (Dong et al. 2019), and *methylGSA* (Ren and Kuan 2019), have been proposed for gene set enrichment analysis (GSEA) of methylation array data. *MethylGSA* is an R Bioconductor package that contains several different gene set testing approaches: mRRA, which adjusts multiple p-values for each gene by Robust Rank Aggregation followed by either over-representation analysis (ORA) or functional class scoring in combination with GSEA, and mGLM, which is an extension of GOglm, implementing a logistic regression to adjust for the number of probes in the enrichment analysis (Mi et al. 2012). The ebGSEA method uses a global test to rank genes, instead of CpGs, based on their total level of differential methylation; enrichment of gene sets is then calculated from the ranked gene list using either a Wilcoxon Test (WT) or Known Population Median Test (KPMT) (Dong et al. 2019). Both ebGSEA and the *methylGSA* methods use individual CpG probe-based differential methylation features, and we are presently not aware of any methods for performing gene set testing for differentially methylated regions.

Here we present GOmeth and GOregion to perform gene set analysis in the context of DNA methylation array data for differential methylation of CpG sites and regions, respectively. The key aspect of our methods is the ability to take into account biases inherent in the data, which relate to how differentially methylated probes are annotated to differentially methylated genes, that can then be assigned to a gene set. Specifically, measured CpG sites are not distributed evenly across the genome, and we and others show that genes that have more CpG sites measured across them are more likely to be detected as differentially methylated compared to genes that have fewer measured CpG sites. Our original implementation of GOmeth addressed this bias (Phipson, Maksimovic, and Oshlack 2016). In addition, approximately 10% of gene-annotated CpGs are assigned to more than one gene, violating assumptions of independently measured genes. Our new and improved implementation of GOmeth considers both of these biases in our approach for detecting enriched pathways. Further to this probe-wise analysis, we developed the GOregion method, which takes these biases into account when performing gene set testing following a region-based analysis. Finally, we have implemented new functionality allowing users to restrict gene set testing to only the probes annotated to specific genomic features, such as probes in promoters of genes.

In this paper we have evaluated the performance of our methods on real and simulated data, as well as comparing to other available methods across a variety of datasets. We found that our methods were the best statistical and computational performers across a variety of comparisons and gene set testing collections. Our methods are publicly available in the Bioconductor R package, *missMethyl*. All of the analysis performed in this paper can be found at the following website: http://oshlacklab.com/methyl-geneset-testing/. The GitHub repository associated with the analysis website is at: https://github.com/Oshlack/methyl-geneset-testing.

## Results

### Composition biases of 450K and EPIC arrays

Consider the scenario where we have performed differential methylation analysis on individual CpG sites. In order to perform gene set enrichment analysis based on the results from a probe-wise differential methylation analysis, we need to annotate each probe on the array to a gene. One approach for gene set testing is to simply call a gene differentially methylated if at least one CpG site associated with that gene is significantly differentially methylated, and this has been used in many previous analyses (e.g. Zhang et al. (2013), Phipson and Oshlack (2014)). The problem with this approach is that the numbers of CpG sites measured across each gene varies significantly across the genome (Figure 1A, Supplementary Figure 1A). For the 450K array, the minimum number of CpGs measured per gene is 1 and the maximum is 1299 with a median of 15, based on the *IlluminaHumanMethylation450kanno*.*ilmn12*.*hg19* annotation package. For the EPIC array, the numbers of CpGs measured across genes ranges from 1 to 1485 (median = 20, *IlluminaHumanMethylationEPICanno*.*ilm10b4*.*hg19* annotation package). Genes that have larger numbers of CpGs measured are more likely to be called differentially methylated when comparing B-cells vs natural killer cells (Figure 1B), and this holds true for the majority of data sets. This bias towards genes with more measured CpG sites can in turn influence the probability of a gene set being called significantly enriched, as some gene sets contain genes with more than the average number of CpGs, and some have genes with fewer measured CpGs (Figure 1C). In this paper, we refer to this particular source of bias as “probe-number bias”.

**Figure 1.**
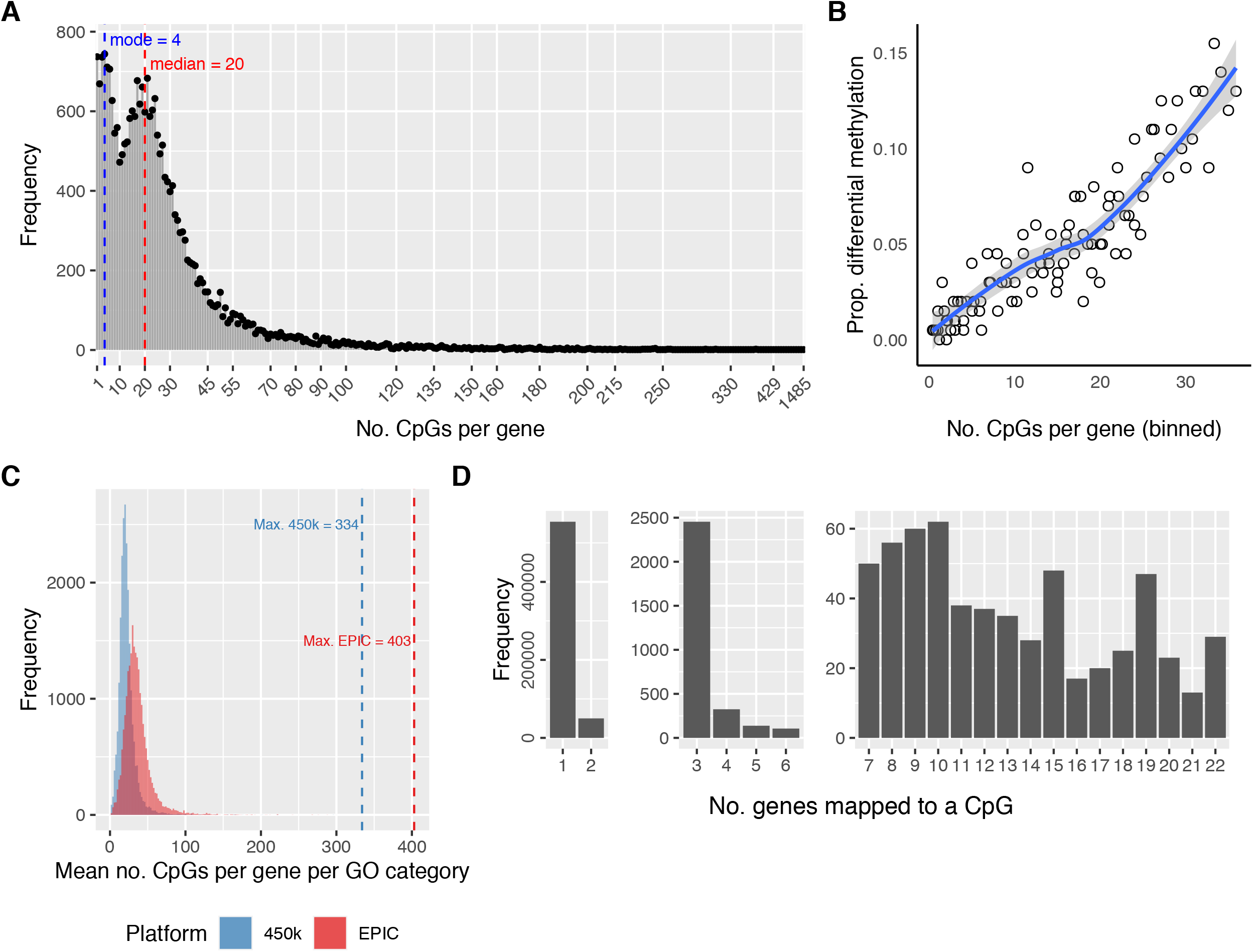
Array design bias for the Illumina HumanMethylation EPIC BeadChip. **(A)** Frequency plot of the numbers of CpGs measuring methylation across each gene for the EPIC array (probe number bias). The most extreme value is 1485 CpGs measuring methylation across a single gene. The median is 20 and the first mode is 4. **(B)** Plot demonstrating probe-number bias for B-cells vs. NK cells from sorted blood cell type EPIC array data. Genes with more measured CpGs are more likely to be differentially methylated. **(C)** Histogram of the median numbers of CpGs per gene for each GO category for the 450K and EPIC arrays. The distributions differ between the arrays, however, both show a varying number of CpGs per gene per GO category. GO categories with more CpGs per gene, on average, have greater power to be significantly enriched. **(D)** Split bar chart showing the numbers of genes annotated to each CpG (multi-gene bias). While the majority of CpGs are annotated to only one gene, there is still a large number annotated to 2 or more genes.

Approximately 70% of the probes on the EPIC array are annotated to at least one gene (74% for 450K array). However, another annotation issue, which is more subtle, is that a single CpG may be annotated to more than one gene as the gene regions overlap on the genome. While the majority of CpGs with gene annotations are associated with only one gene (329,365/359,832 = 92% for 450K, 554,221/607,820 = 91% for EPIC arrays), there are still a large number of CpGs annotated to 2 or more genes (Figure 1D, Supplementary Figure 1B). This can cause issues as the measurements of differentially methylated genes are not independent. If we use every gene associated with a single CpG, we risk counting a single significant CpG site multiple times when including the genes as enriched in a gene set of interest. If these genes were evenly distributed across the GO categories, this may not be an issue. However, genes that are close in genomic proximity can be functionally related and be present in a single GO category. An extreme example of this is cg17108383 which is annotated to 22 genes, all belonging to the protocadherin gamma gene cluster (Supplementary Figure 2). All 22 of these genes are present in the GO category “GO:0007156: homophilic cell adhesion via plasma membrane adhesion molecules”, which contains a total of 129 genes. If each of these significant genes are included when performing a hypergeometric test for enrichment of the GO category, then for this single significant CpG site, the overlap between the differentially methylated genes and the genes in the gene set is increased by 22, and this GO category will appear significantly enriched. Unfortunately, this is not an isolated occurrence. For the EPIC array, gene ontology (GO) enrichment analysis on the 53,599 CpGs that are annotated to at least 2 genes results in 114 significantly enriched GO categories (Holm’s adjusted p-value cut-off < 0.05). Restricting to CpGs that are annotated to at least 3 genes results in significant enrichment of 56 GO categories (Holm’s adjusted p-value cut-off < 0.05), which are mostly related to processes involved in transcriptional regulation (Supplementary Table 1).

We refer to this newly identified source of bias as “multi-gene bias”. In order to reduce false positives, it is important to take this multi-gene bias into account when calculating the intersection between differentially methylated genes and the genes in each gene set. One approach for dealing with multi-gene bias is to simply randomly select one gene to be represented by the CpG, but this approach risks losing valuable information by ignoring the remaining associated genes. We include this multi-gene bias in our statistical framework for gene set testing using a weighting strategy (Figure 2) and ensure significant CpGs are only counted once, at most.

**Figure 2.**
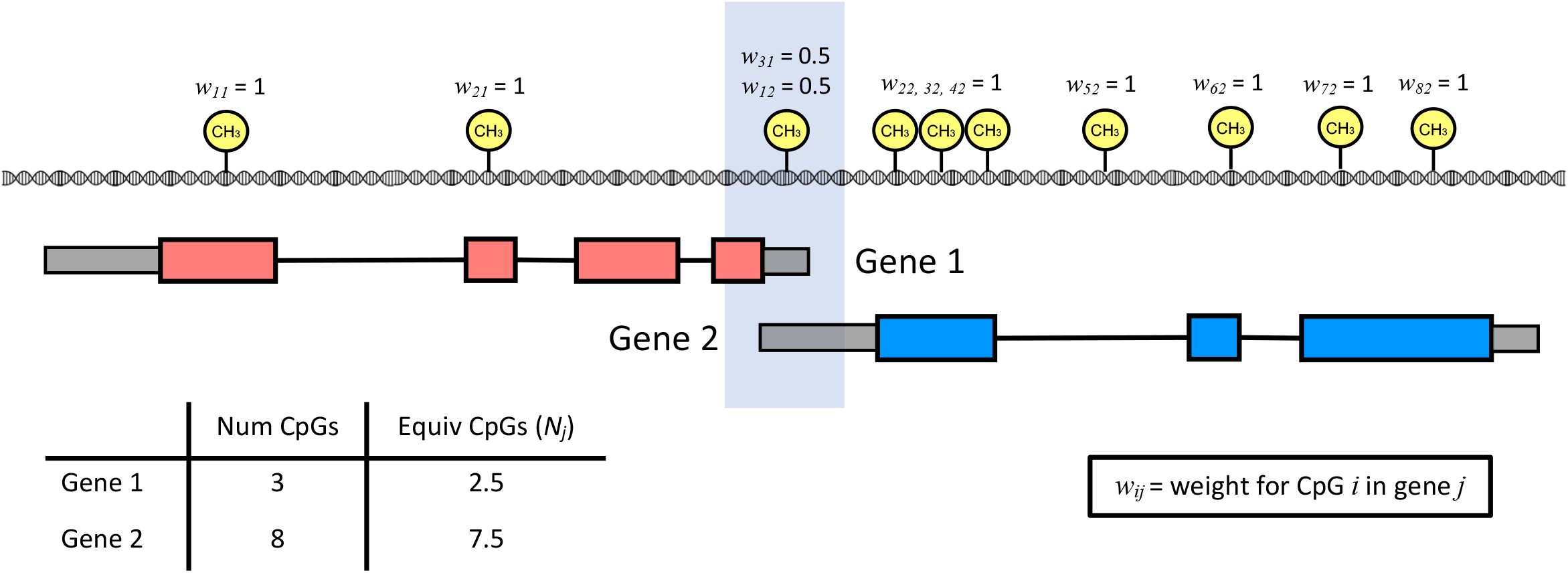
Overview of how probe number and multi-gene bias is taken into account in GOmeth. CpGs are not evenly spaced throughout the genome. Gene 1 has methylation measured at three CpGs and gene 2 has methylation measured at eight CpGs. The CpG shaded in blue is an example of a shared genomic location between gene 1 and 2, with one CpG measuring the methylation status for two genes and is thus not independently measured. By calculating a weight for each CpG inversely proportional to how many genes that CpG is annotated to, we can calculate the equivalent numbers of CpGs measured across each gene and ensure the enrichment statistic for Wallenius’ noncentral hypergeometric test is not artificially inflated due to multi-gene annotated CpGs.

### GOmeth performs gene set testing on differentially methylated CpG sites

Our method for gene set testing performs enrichment analysis of gene sets while correcting for both probe-number and multi-gene bias in methylation array data. The detailed statistical approach is outlined in the ‘Methods’ section. This method was inspired by GOSeq (Young et al. 2010). The GOSeq method was designed to account for the fact that longer genes have more sequencing read counts compared to shorter genes, and hence have more power to be statistically significantly differentially expressed. Similarly, we see that genes with a larger number of measured CpGs have a higher probability of differential methylation (Figure 1B). To account for this bias, GOmeth empirically estimates the probability of differential methylation of a gene as a function of the number of CpGs. In generating the trend, we adjust the number of CpGs in a gene to account for those that are assigned to multiple genes by using a fractional count. As the probability of differential methylation is calculated empirically from the data, this trend may look different between datasets, but generally we always observe a strong positive trend. From the empirical trend, the odds of differential methylation for each gene set or gene ontology category is calculated based on the average probabilities of genes in the set. The intersection of the number of differentially methylated genes in the gene set of size *g* also needs to be defined. Rather than just assigning all genes with a differentially methylated CpG equally, we also take into account the CpGs annotated to multiple genes in this step. Specifically, we assign a weight, *w*_*ij*_, to each CpG that is the reciprocal of the number of genes annotated to the CpG (Figure 2). We then sum all the CpG weights assigned to a gene and add one to the overlap if the weights are greater than one and sum(*w*_*ij*_) otherwise. If all CpGs were annotated to exactly one gene, then the weight, *w*_*ij*_, would be one for every CpG, and the overlap would be the number of genes with a significant CpG. However, if a significantly differentially methylated CpG is annotated to 2 genes, the CpG is assigned a weight of 0.5 to each gene. If no other significant CpGs are annotated to that particular gene, then the gene will contribute a “count” of 0.5 to the intersection statistic for the Wallenius’ noncentral hypergeometric test. Thus, if both genes are present in the same gene ontology category, they will contribute a total count of at most one to the intersection statistic. For genes with multiple significant CpGs that may include several multi-gene associated CpGs, the maximum total count a gene can contribute to the intersection statistic is 1.

Next, a test based on Wallenius’ noncentral hypergeometric distribution is performed for each gene set or gene ontology category, incorporating the odds that a gene set is more or less likely to be enriched based on the probe-number bias of the experiment with the intersection statistic defined above (Phipson, Maksimovic, and Oshlack 2016).

We have implemented our approach to gene set testing with two different functions in the *missMethyl* Bioconductor R package, ‘gometh’ and ‘gsameth’. The difference between the two functions is minimal, with ‘gometh’ specifically testing for enrichment of gene ontology (GO) categories from the *GO*.*db* annotation package, or KEGG pathways from the *KEGG*.*db* annotation package. The ‘gsameth’ function is a more general version of ‘gometh’, where the user can supply any list of gene sets to be tested. In addition, the ‘gometh’ and ‘gsameth’ functions allow the set of significantly differentially methylated CpGs to be restricted to genomic regions of interest such as promoters or gene bodies, as these may interrogate different biological pathways.

### Improved Type I error rate control with GOmeth

We first tested the performance of GOmeth by randomly sampling sets of CpG probes from the EPIC and 450K array annotation that we designated as differentially methylated and running gene ontology analyses. Under these null scenarios we would not expect to see significant enrichment of any GO categories. We randomly selected 100 sets each of 50, 100, 500, 1000, 5000 and 10,000 CpGs as “significantly” differentially methylated based on the 450K and EPIC annotation. We tested for enrichment of gene ontology sets for each simulation and calculated the number of GO categories that were significant at a p-value threshold of 0.05. For a test to correctly control the type I error rate, we expect 5% or fewer GO categories to have significant p-values for random data. We compared the three testing options available in GOmeth: the hypergeometric test with no bias corrections (“HGT”), Wallenius’ noncentral hypergeometric test taking into account probe-number bias only (“HGT-mod”) and GOmeth, which takes into account both probe-number and multi-gene bias (Figure 3A & B, Supplementary Figure 3A & B). Under these simulation conditions we were unable to compare to the *methylGSA* methods and ebGSEA since these tests require M-values or βvalues as input rather than just the list of significant probes.

**Figure 3.**
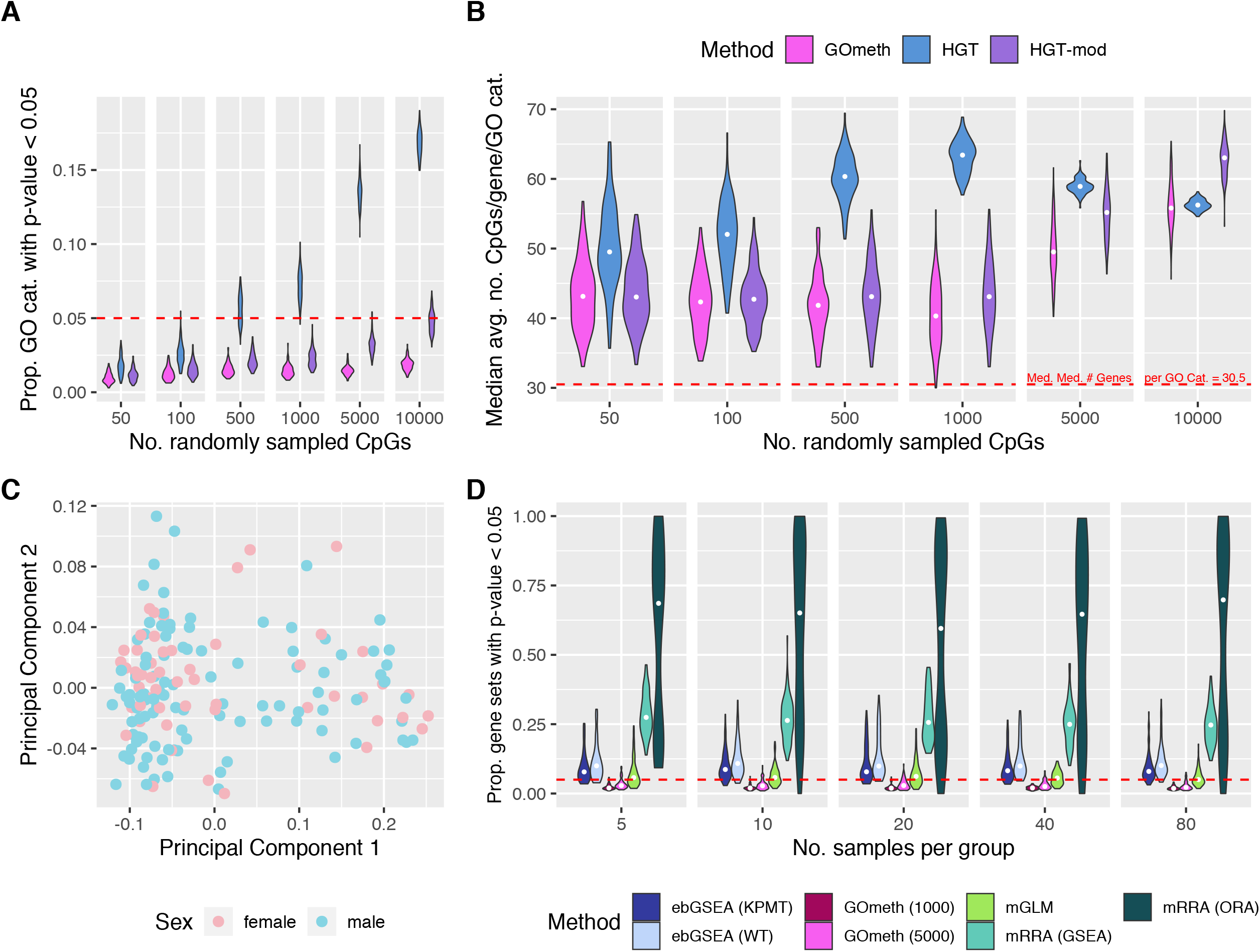
Evaluation of false discovery rate control for EPIC array data. **(A)** Type I error rates across 100 simulations for varying numbers of randomly sampled CpGs. **(B)** Median average numbers of CpGs per gene for GO categories with an unadjusted p-value < 0.05. The hypergeometric test is biased towards GO categories with more CpGs per gene on average. GOmeth = adjust for probe-number and multi-gene bias; HGT = hypergeometric test; HGT-mod = adjust for probe-number bias only. **(C)** Multidimensional scaling plot of normal samples from TCGA KIRC data, coloured by sex. **(D)** False discovery rate control of seven gene set testing methods using normal samples from TCGA KIRC data. Two groups were generated by randomly sampling *n* samples per group, followed by differential methylation analysis and subsequent gene set testing. This was repeated 100 times at each sample size. The proportion of gene sets with unadjusted p-value < 0.05 across the 100 null simulations is shown for each method, at each sample size. Methods with good false discovery rate control should have relatively tight distributions around the red dashed line at 0.05. ebGSEA (KPMT) = ebGSEA using Known Population Median Test; ebGSEA (WT) = ebGSEA using Wilcoxon Test; GOmeth (1000) = GOmeth using top 1000 ranked probes; GOmeth (5000) = GOmeth using top 5000 ranked probes; mGLM = methylglm; mRRA (GSEA) = methylRRA using gene set enrichment analysis; mRRA (ORA) = methylRRA using over-representation analysis.

Figure 3A summarises the type I error rates for our methods across the varying sets of randomly selected CpGs sampled from the EPIC array. As the numbers of differentially methylated CpGs increased we noted that the hypergeometric test reported too many significant GO categories, particularly for more than 500 CpGs. Clearly, taking into account the probe-number bias makes the biggest correction to false discoveries, with HGT-mod and GOmeth maintaining the correct Type I error rate. Correcting for multi-gene bias (GOmeth) further reduced the numbers of significantly enriched GO categories. We did notice that not accounting for the multi-gene bias led to an increase in false discoveries as the numbers of significant CpGs increased. A similar trend was observed for simulations based on the 450K array (Supplementary Figure 3A).

Of the significant GO categories reported by each method, we found that the hypergeometric test was more biased towards GO categories with more CpGs measured per gene on average (Figure 3B, Supplementary Figure 3B). HGT-mod and GOmeth reported significant categories that had a wide range of CpGs per gene on average. Based on the results of these simulations, we selected GOmeth taking into account both probe-number and multi-gene bias as the best option to use for further analysis, even though it is quite a conservative test in this particular simulation scenario.

The prior simulations are limited in that they only generate a random set of CpGs to test for over-representation of gene sets. However, previously published methods, ebGSEA and the *methylGSA* methods, require differential methylation measurements (M-values or βvalues) as input. In order to compare the Type I error rate control of GOmeth with ebGSEA and *methylGSA* we analysed the normal samples from the TCGA 450K array kidney renal cell carcinoma (KIRC) dataset (Figure 3C), as well as the normal samples from the TCGA 450K breast invasive carcinoma (BRCA) dataset (Supplementary Figure 3C). For both datasets, we took a resampling approach whereby we randomly assigned samples to one of two “groups” and varied the sample size per group (n = 5, 10, 20, 40, 80). We then performed differential methylation analysis between the two artificial groups, followed by gene set testing using the available methods: mGLM, mRRA (ORA), mRRA (GSEA), ebGSEA (WT), ebGSEA (KPMT), GOmeth with top 1000 CpGs and GOmeth with top 5000 CpGs. While the ebGSEA methods are available in the *ChAMP* R Bioconductor package, we used a more flexible implementation available in the *ebGSEA* R package on GitHub. We defined the input for GOmeth as either the top 1000 or top 5000 most highly ranked differentially methylated CpGs even though the probes did not reach statistical significance. This allowed us to calculate the proportions of gene sets that were significantly enriched for each of the methods, where “significant” is defined as a p-value less than 0.05. We repeated these steps 100 times at each sample size. In this scenario, where there are no true biological pathways differentially methylated, we expect 5% or fewer gene sets to be significantly enriched. We used the 8567 gene sets from the Broad’s Molecular Signatures Database (MSigDB) available in the *ChAMP* package to evaluate the false discovery rate. In addition, to ensure that the methods in the *methylGSA* package produced reasonable output, we limited the size of the gene sets to those with at least 5 and at most 5000 genes in the set. If these additional constraints are not included, the *methylGSA* methods mGLM and mRRA (ORA) produced results that were heavily biased towards reporting very small gene sets as highly significant. Furthermore, mRRA (ORA) was also biased towards ranking large gene sets very highly, if they were not filtered out (Supplementary Figure 3E).

In total, we compared seven different variants of the gene set tests: GOmeth with top 1000 CpGs, GOmeth with top 5000 CpGs, the three testing frameworks in *methylGSA* and the two tests in ebGSEA. In general, varying the sample size did not make a difference to the results, with consistent patterns observed for all sample sizes, across the two datasets (Figure 3D, Supplementary Figure 3D). The worst performing test was mRRA (ORA), which had a median proportion of significantly enriched gene sets of at least 0.6 (or a median of 5388 enriched gene sets, across the 5 sample sizes). mRRA (GSEA) also performed poorly, with a median proportion of significantly enriched gene sets of at least 0.25 (or a median of 2117 enriched gene sets, across the 5 sample sizes). However, mGLM correctly controlled the false discovery rate at 0.05. The two variants of ebGSEA had slightly greater than 0.05 median proportion of false discoveries although their results were more variable across the 100 simulations at each sample size, with some simulations showing large numbers of false positives. GOmeth, using both top 1000 and 5000 CpGs, showed highly consistent performance, with median proportions of false discoveries < 0.05 (with a median of 180 and 278.5 enriched gene sets, respectively, across the 5 sample sizes), suggesting that GOmeth correctly controls for false discoveries. The performance of the seven methods were highly consistent across both the KIRC and BRCA datasets.

### Application to blood cell type EPIC data

Following our simulation studies, we wanted to test the performance of GOmeth, ebGSEA and the *methylGSA* methods on real data that contained differential methylation. We used a publicly available dataset of flow sorted blood cell types profiled on Illumina Infinium HumanMethylationEPIC arrays (GSE110554) (Salas et al. 2018). Cell types are easily distinguished based on methylation patterns (Figure 4A). We chose to perform our differential analysis and gene set testing on three independent pair-wise comparisons of cell types with varying numbers of differentially methylated probes: (1) CD4 vs CD8 T-cells, (2) monocytes vs neutrophils and (3) B-cells vs natural killer (NK) cells (Figure 4B). Differential methylation was performed using TREAT (McCarthy and Smyth 2009) and CpGs were defined as significantly differentially methylated if they had false discovery rates < 0.05 and Δβcut-off of ∼ 10% (corresponding to ΔM ∼ 0.5). Following the differential methylation analysis, we tested enrichment of GO sets and KEGG pathways using hypergeometric tests, GOmeth, the three *methylGSA* methods and the two ebGSEA tests. Again, we limited the gene sets to those with a minimum of 5 and a maximum of 5000 genes for *methylGSA*. For input to GOmeth, we used either the statistically significant CpGs at FDR < 0.05 (CD4 vs CD8 T-cells) or the top 5000 most highly ranked CpGs for comparisons with more than 5000 significant CpGs (B-cells vs NK cells and monocytes vs neutrophils).

**Figure 4.**
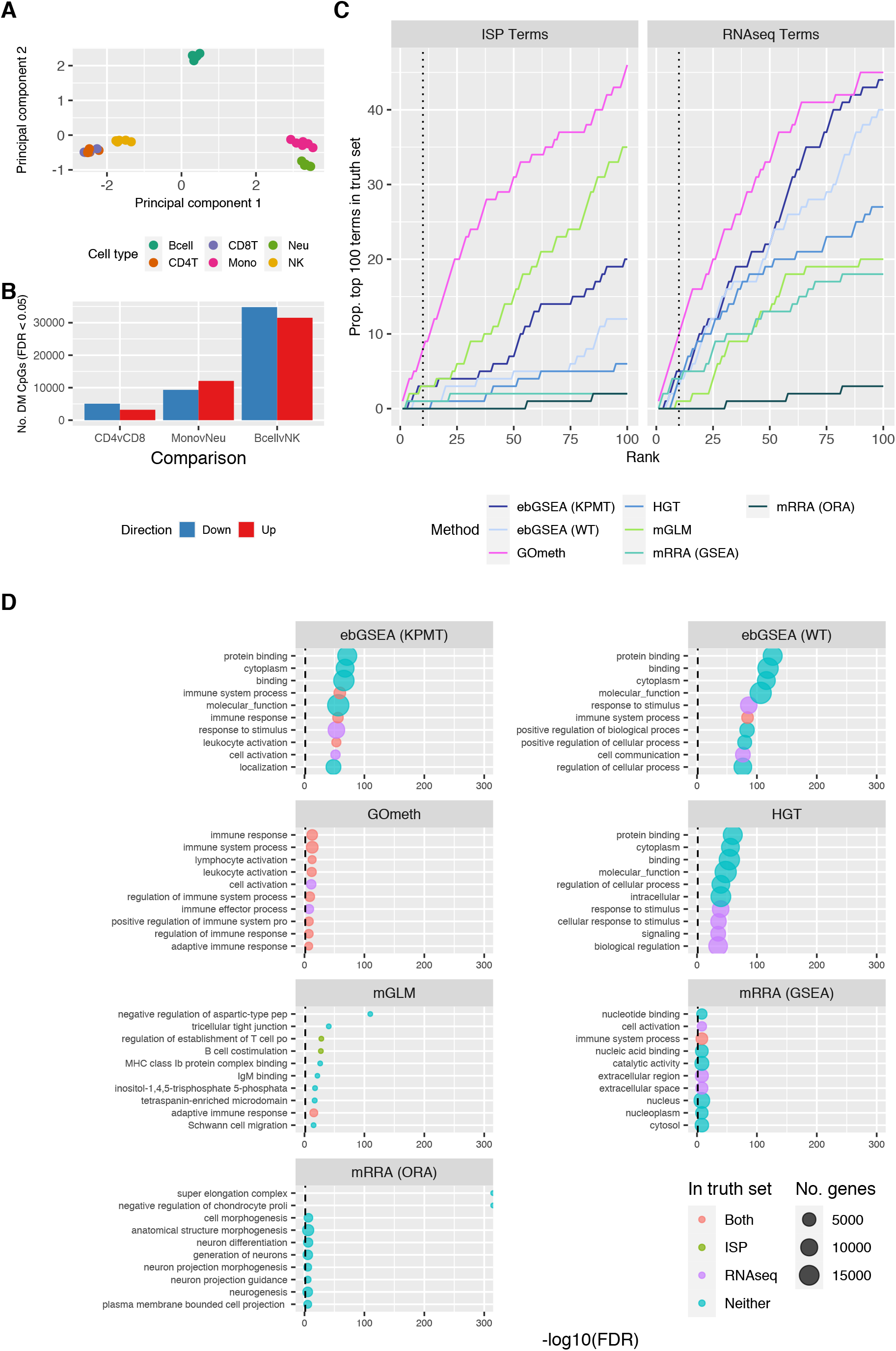
Comparison of gene set testing performance on Gene Ontology (GO) categories. **(A)** Multidimensional scaling plot of EPIC array sorted blood cell data. **(B)** Numbers of differentially methylated CpGs with an adjusted p-value < 0.05, for each cell type comparison: CD4 T-cells vs. CD8 T-cells, monocytes vs. neutrophils and B-cells vs. NK cells. The blue bar is the number of significant CpGs that are less methylated in the first cell type relative to the second, and the red bar is the number that are more methylated; e.g. ∼5000 CpGs are less methylated and ∼3000 are more methylated in CD4 T-cells, compared to CD8 T-cells. **(C)** Cumulative number of GO terms, as ranked by various methods, that are present in each truth set for the **B-cells vs. NK** comparison. ISP Terms = immune-system process child terms truth set; RNAseq Terms = top 100 terms from RNAseq analysis of the same cell types. **(D)** Bubble plots of the top 10 GO terms as ranked by various gene set testing methods. The size of the bubble indicates the relative number of genes in the set. The colour of the bubble indicates whether the term is present in either RNAseq (purple) or ISP (green) truth sets, both (red) or neither (blue). ebGSEA (KPMT) = ebGSEA using Known Population Median Test; ebGSEA (WT) = ebGSEA using Wilcoxon Test; GOmeth = GOmeth using either FDR < 0.05 (for contrasts with <5000 significant CpGs) or top 5000 most significant probes; HGT = hypergeometric test; mGLM = methylglm; mRRA (GSEA) = methylRRA using gene set enrichment analysis; mRRA (ORA) = methylRRA using over-representation analysis.

To evaluate the significant gene sets in a systematic way we took two approaches. First, we identified all immune categories in the GO database, as these are expected to be highly enriched when comparing different blood cell types. We therefore defined “true positive” GO categories as all the child terms under the parent GO category “immune system response” (GO:002376) from AMIGO 2 (http://amigo.geneontology.org/amigo/term/GO:0002376). We then counted how many of these immune sets were present in the top ranked gene sets for each method. We similarly defined true positives for the KEGG pathways by identifying all pathways belonging to the following categories: Immune system, Immune disease, Signal transduction, Signaling molecules and interaction (https://www.genome.jp/kegg/pathway.html). The second approach we took to evaluate the different methods was by analysing a publicly available RNA-Seq dataset comparing the same sorted blood cell types (GSE107011; SRP125125) (Monaco et al. 2019; W. Xu et al. 2019). We performed differential expression analysis and gene set testing on the expression data and defined the top 100 significantly enriched gene sets from the RNA-Seq analysis as the “truth” (Supplementary Figure 4-D).

For GO categories, GOmeth consistently performed the best with the highest numbers of top ranked categories overlapping with “truth” sets across the three comparisons (Figure 4C-D, Supplementary figures 5-). For KEGG pathways, the differences between the methods was not as clear, with mGLM, ebGSEA and GOmeth generally performing well. The mRRA (GSEA) and mRRA (ORA) methods had highly variable performance (Supplementary Figure 7). For the monocyte vs neutrophil comparison, the ebGSEA methods had more power to detect significant gene sets (Supplementary figure 7E).

Next we examined the top 10 ranked terms for each of the seven gene set testing methods (Figure 4D, Supplementary figures 5-7). For the B-cells vs NK cells, we noted that the hypergeometric test, and to a lesser extent, the two ebGSEA methods, tended to have very large, non-specific GO categories most highly ranked, with “protein binding”, “cytoplasm” and “molecular function” in the top 10 (Figure 4D). The top 10 enriched GO categories for GOmeth were more biologically relevant, with immune specific gene set tests highly enriched (for example “leukocyte activation” and “lymphocyte activation”). All of the top 10 terms for GOmeth were included in at least one “truth” set. For the *methylGSA* methods, mGLM appeared to have more immune specific categories in the top 10 (e.g. “adaptive immune response” and “ regulation of leukocyte cell-cell adhesion”) compared to mRRA (GSEA) and mRRA (ORA). The results for mRRA (ORA) and mRRA (GSEA) were more difficult to interpret with none of the top 10 mRRA (ORA) results included in either “truth” set. A similar pattern was observed for the CD4 vs CD8 T-cells and monocyte vs neutrophil comparisons, with GOmeth consistently ranking more immune specific terms in the top 10 (Supplementary figures 5B and 6B).

### Application to B-cell development 450K data

We applied the seven different gene set testing methods to a developing pre B-cell dataset which measured methylation using the 450K array, with matched gene expression captured using Affymetrix arrays. Four populations of B-cell developmental stages were isolated from human fetal bone marrow, from 8 individuals. We focused on the comparison between Stage 1 (multipotent progenitor cells) and Stage 2 (pre B-cells I) and analysed the methylation data taking a similar approach as described above (Supplementary Figure 8A, B). For the gene expression dataset, background correction and normalisation was performed using the RMA algorithm (Bolstad et al. 2003; Irizarry et al. 2003), low intensity probes were filtered out and differential expression analysis was performed with *limma* (Supplementary Figure 4E-G). We created the gene expression “truth” sets by selecting the top 100 gene sets for GO and KEGG pathways identified using the ‘goanna’ and ‘kegga’ functions in *limma*. We also used the immune related sets from GO and KEGG as previously described, as alternative “truth” sets.

For GO categories, GOmeth was the top performer, with ebGSEA (KPMT) performing consistently well across both “truth” sets (Supplementary Figure 8C, D). The *methylGSA* methods did not perform as well on this dataset, with mGLM the best of the three. For KEGG pathways, all methods performed similarly (Supplementary Figure 9A), however the gene sets were not very highly statistically significant (Supplementary Figure 9B). For the top 10 ranked GO categories, GOmeth was the only method that had all 10 present in either “truth” set (Supplementary Figure 8D). For KEGG pathways, 8/10 pathways for GOmeth were present in either “truth” set and the top ranked pathway for GOmeth was “Hematopoietic cell lineage” (Supplementary Figure 9B).

We generally found, across all dataset comparisons and gene set ensembles, that the hypergeometric test, mRRA (ORA) and mRRA (GSEA) tended to rank large, non-specific categories very highly, with GOmeth ranking more biologically relevant gene sets in the top For the two ebGSEA methods, ebGSEA (KPMT) appeared to perform better than ebGSEA (WT), however for dataset comparisons with a large number of significant probes, the ebGSEA methods tended to rank large, non-specific categories more highly. For comparisons with fewer significant probes, ebGSEA (KPMT) showed good performance with more power to detect significant enrichment of gene sets. The mGLM method was the best performer of the three methods available from *methylGSA*.

### Comparing compute time between gene set testing methods

While mGLM generally performs well, computationally, it was one of the slowest methods to run on a single core (∼48 minutes), with ebGSEA also taking approximately 50 minutes to complete an analysis (Table 1). The run time of ebGSEA and mGLM can be significantly reduced by parallelising the computations. Using 9 cores, ebGSEA took approximately 25 minutes, and mGLM took just under 20 minutes, to complete the analysis of 8567 MSigDB sets. By comparison, GOmeth is ∼90 times faster than mGLM and ebGSEA. mRRA (ORA) is the fastest to run but generally does not perform as well as other methods.

**Table 1:** Average run-time across all contrasts. Gene sets used are Broad MSigDB gene sets from the *ChAMP* package. All methods were run on a single core.

| Method | Minutes |
| --- | --- |
| mRRA (ORA) | 0.13 |
| GOMeth | 0.54 |
| mRRA (GSEA) | 3.04 |
| mGLM | 47.95 |
| ebGSEA | 50.12 |

### Gene set testing following a region based analysis

CpGs are not evenly spaced across the genome and often appear in clusters e.g. CpG islands (Gardiner-Garden and Frommer 1987); and several studies have demonstrated that CpGs in close proximity have correlated methylation levels (Eckhardt et al. 2006). Thus, rather than testing individual CpGs, identifying correlated methylation patterns between several spatially adjacent CpGs has been shown to yield more functionally relevant results (Hansen et al. 2011).

Several tools have been published for identifying differentially methylated regions (DMRs) from methylation array data: *Probe lasso* (Butcher and Beck 2015), *bumphunter* (Jaffe et al. *2012), DMRcate* (Peters et al. 2015), *mCSEA* (Martorell-Marugán, González-Rumayor, and Carmona-Sáez 2019) and *DMRforPairs* (Rijlaarsdam et al. 2014). Depending on the input data, region finding tools can identify several hundred or even thousands of DMRs. They all generally output the location of the region including the chromosome, region start and region end positions, along with some additional metrics and statistical significance. Some tools attempt to annotate the regions with genes but others do not. Thus, when faced with a long list of DMRs it is unclear how to interpret the biological significance of the results, and there are no gene set testing tools available for DMRs.

To address this, we have developed GOregion; an extension of GOmeth, that enables gene set testing of DMRs. The “goregion” function tests GO terms and KEGG pathways, whilst “gsaregion” is a generalised function that accepts any list of gene sets as input. We reasoned that because region detection is inherently dependent on CpG probe density, DMRs are more likely to be identified in genes with more CpG probes. This trend is observed in the blood cell type and B-cell development data (Figure 5A, Supplementary figure 10A, 11A, 14A). To take this bias into account, GOregion utilises the GOmeth testing framework. GOregion accepts a ranged object of DMRs that have been identified by the user’s choice of region-finding software. These regions are then overlapped with the locations of the CpGs on the Illumina array to identify a set of CpGs underlying the DMRs. These CpG probes are then passed to GOmeth and GOmeth’s existing algorithm is used to test for enrichment of gene sets.

### GOregion applied to sorted blood cell type EPIC data

Using the sorted blood cell EPIC data we identified DMRs for the same three cell type comparisons using the *DMRcate* package: 1) CD4 vs CD8 T-cells, 2) monocytes vs neutrophils and 3) B-cells vs NK cells. Using default parameters, *DMRcate* identified 6404, 7176 and 23,210 differentially methylated regions, respectively. We further filtered the DMRs by only including regions containing 3 or more CpGs with an absolute mean βvalue difference greater than 0.1. This left 789, 1633 and 4723 DMRs, respectively, for downstream analysis (Figure 5B). We then performed gene set testing of GO categories using GOregion, and compared it to a simple approach of overlapping DMRs with known genes, and then testing using a HGT.

**Figure 5.**
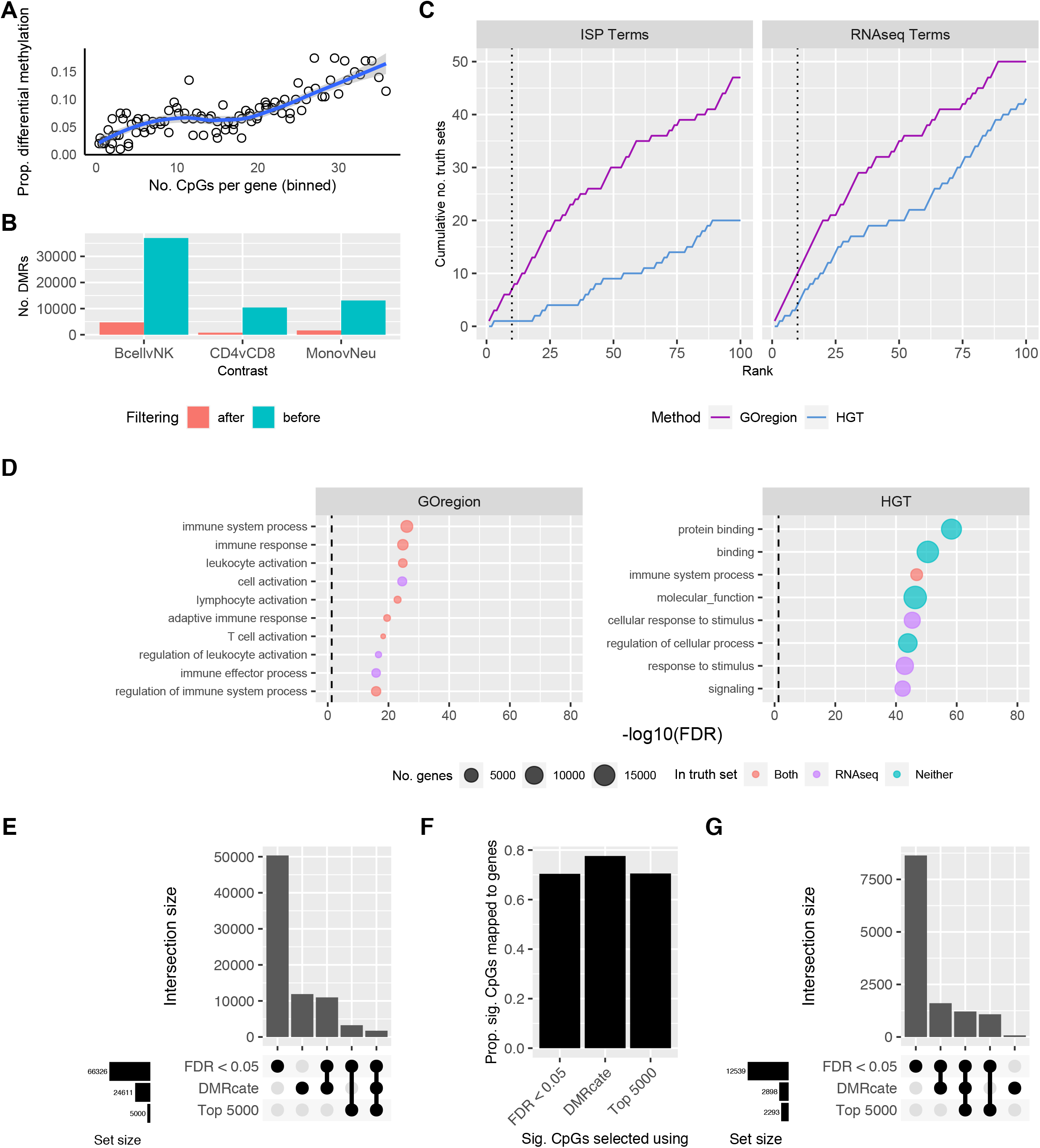
Evaluation of the performance of GOregion on sorted blood cell data. **(A)** Bias plot showing that genes with more measured CpGs are more likely to have a differentially methylated region (DMR). This plot is produced from EPIC array sorted blood cell type data, comparing B-cells to NK cells. **(B)** Numbers of DMRs identified by *DMRcate*, for each cell type comparison: CD4 T-cells vs. CD8 T-cells, monocytes vs. neutrophils and B-cells vs. NK cells. The blue bar is the number of DMRs before filtering, the pink bar is the number of DMRs after filtering out DMRs with < 3 underlying CpGs and an absolute mean |Δβ| < 0.1. **(C)** Cumulative number of GO terms, as ranked by GOregion and a simple hypergeometric test (HGT), that are present in each truth set for the B-cells vs. NK comparison. ISP Terms = immune-system process child terms truth set; RNAseq Terms = top 100 terms from RNAseq analysis of the same cell types. **(D)** Bubble plots of the top 10 GO terms as ranked by GOregion and a simple HGT for the B-cells vs. NK comparison. The size of the bubble indicates the relative number of genes in the set. The colour of the bubble indicates whether the term is present in either RNAseq (purple) or ISP (green) truth sets, both (red) or neither (blue). **(E)** Upset plot showing the characteristics of the CpGs selected as “significant” for the B-cell vs. NK comparison by a probe-wise differential methylation analysis using a significance cut off (FDR < 0.05), the top 5000 CpGs as ranked by the probe-wise analysis (Top 5000) or the CpGs underlying the filtered *DMRcate* regions (DMRcate). The probe-wise analysis with FDR < 0.05 identified over 60,000 CpGs as “significant” and had the most unique CpGs. However, despite identifying fewer “significant” CpGs (∼25,000), almost half of the CpGs identified by DMRcate are unique (∼12,000). **(F)** Proportion of “significant” CpGs that are annotated to genes as identified by the three different strategies. **(G)** Upset plot showing the characteristics of the genes that “significant” CpGs are annotated to, as identified by the three different strategies, for the B-cell vs. NK comparison. CpGs identified by the probe-wise analysis with FDR < 0.05 map to over 12,000 genes. Although the CpGs identified by DMRcate map to far fewer genes (∼2900), a number of them are unique to this approach.

As previously described, we evaluated the results by counting the numbers of highly ranked immune-related GO terms, and the numbers of highly ranked GO categories identified in the RNA-Seq data analysis of the same cell types. GOregion consistently ranked immune-related and RNA-Seq “truth” terms more highly than the simple HGT-based strategy (Figure 5C, Supplementary figures 10B, 11B). Examining the top 10 most highly ranked GO categories showed that GOregion categories are more specific to immune processes than those identified using the HGT approach (Figure 5D, Supplementary figure 10C, 11C). Across all comparisons, GOregion always ranked more “truth” sets in the top 10 than HGT. For example, the top 10 gene sets ranked by GOregion for the B-cells vs NK comparison were highly specific to immune system processes, e.g. “T cell activation”, whereas for HGT the sets were very broad and contained thousands of genes e.g. “protein binding” (Figure 5D).

We wanted to explore the differences between a region analysis and a probe-wise analysis and how this would affect gene set testing results. GOmeth is highly dependent on the set of significant CpGs provided as input, so we postulated that selecting CpGs using a region-level analysis could be more biologically relevant, in certain circumstances. We compared the numbers of differentially methylated probes and genes that are selected based on a region analysis with *DMRCate*, a probe-wise analysis using an FDR cut-off < 0.05, and a probe-wise analysis selecting the top 5000 differentially methylated CpGs. For the B-cell vs NK cells comparison, a probe-wise analysis with FDR < 0.05 selects over 60,000 differentially methylated CpGs. More than 50,000 of these CpGs are unique to this approach and not identified with the region approach, whereas *DMRCate* identifies ∼24,000 CpGs in DMRs (Figure 5E). We noted that although *DMRCate* always captured fewer significant CpGs (Figure 5E, Supplementary figure 10D, 11D), a higher proportion of the total significant CpGs were annotated to genes compared with either of the probe-wise approaches (Figure 5F, Supplementary Figures 10E, 11E). This results in sets of genes uniquely captured using a region-based analysis (Figure 5G, Supplementary Figures 10F, 11F).

Comparing GOregion to the two probe-wise approaches using our previously defined “truth” sets showed that all of the approaches performed similarly well across all the contrasts, except for B-cells vs NK cells (Supplementary figure 12A, B). For that comparison, GOregion and the GOmeth probe-wise analysis using the top 5000 CpGs both performed markedly better than GOmeth with significant CpGs selected using FDR < 0.05 (Supplementary figure 12A). This is likely due to these significant CpGs being annotated to > 12,000 genes, resulting in a highly non-specific set of genes as input to GOmeth, whereas *DMRCate* regions are annotated to just under 3000 genes (Figure 5G). Examining the terms that were highly ranked by the various approaches, across the different contrasts, revealed that they all tended to be immune specific (Supplementary figure 12C-E). The exception is the B-cells vs NK comparison produced by GOmeth based on CpGs selected at FDR < 0.05, where the most highly ranked terms were very broad categories such as “cell communication”, “signalling” and “plasma membrane part” (Supplementary figure 12C). Hence, for comparisons which result in very large numbers of significantly differentially methylated CpGs, limiting the set of CpGs used as input for gene set testing, such as performing a region analysis, is important for producing meaningful results.

Region-finding software is itself dependent on numerous parameters and appropriate downstream filtering of results can also be important in identifying the most biologically informative set of DMRs. Using the blood cell type dataset, we demonstrate that different DMR filters affect the downstream GOregion gene set testing results in different ways, depending on the comparison (Supplementary Figure 13). For our dataset, the most important filter to include is a mean Δβdifference cut-off, i.e. the size of the methylation difference. For this particular dataset, we find that a Δβdifference cut-off of 0.1 performs best across all comparisons. For the B-cells vs NK cells comparison, we noticed that not filtering with a Δβ cut-off produced very poor results. In contrast, for the monocyte vs neutrophils comparison, a Δβ cut-off of 0.2 was too stringent and resulted in fewer gene sets overlapping with the “truth” sets (Supplementary figure 13A, B).

### GOregion applied to B-cell development 450K data

Using *DMRcate*, we identified 2151 DMRs when comparing Stage 1 vs Stage 2 B-cells, and observed a strong probe-number bias trend (Supplementary Figure 14A). Once again, we compared the top ranked GO terms for GOregion and HGT with our two “truth” sets: 1) the top 100 ranked gene sets from comparing Stage 1 vs Stage 2 B-cells in the Affymetrix gene expression data, and 2) the immune related terms. GOregion outperformed HGT, with a greater overlap of the top ranked gene sets with both “truth” sets (Supplementary Figure 14B). Examining the top 10 GO terms showed that GOregion had 7/10 terms present in either “truth” set, with only 3/10 gene sets overlapping either “truth” set for HGT. Two of the top 10 GO terms identified by GOregion, but not identified in either “truth” sets, were related to cytokine production, which is known to be important in B-cell development (Vazquez, Catalan-Dibene, and Zlotnik 2015). HGT ranked very broad, non-specific GO terms very highly (e.g. “molecular function”, “protein binding”), whereas GOregion ranked more immune related gene sets in the top 10 (e.g. “leukocyte activation”).

We compared the numbers of differentially methylated probes and genes selected using either a region-based analysis, a probe-wise analysis with FDR < 0.05 or a probe-wise approach selecting the top 5000 differentially methylated CpGs (Supplementary Figure 14D-F). For Stage 1 vs Stage 2 developing B-cells, 11,597 probes were selected using *DMRCate* compared to only 3148 with a probe-wise analysis with FDR < 0.05, suggesting that thousands of probes that are not individually significant were underlying regions (Supplementary Figure 14D). A higher proportion of probes captured using *DMRcate* were annotated to genes compared to the probe-wise analyses (Supplementary Figure 14E), and although *DMRcate* selected 11,597 probes, these were annotated to only 1,664 genes (Supplementary Figure 14F). As we observed in the sorted blood cell type data, the *DMRcate*-selected CpGs overlapped genes that were unique to the region-based analysis (Supplementary Figure 14F).

For this particular dataset, GOmeth and GOregion performed very similarly with all methods (GOmeth (top 5000), GOmeth (FDR<0.05) and GOregion) showing a similar degree of overlap with the two “truth” sets (Supplementary Figure 15A). While there were some differences in the top 10 ranked GO terms, they were all involved in immune system processes (Supplementary Figure 15B). Although we did not filter the DMRs prior to gene set testing with GOregion for this particular dataset, we investigated the effect of filtering on the overlap with the “truth” sets (Supplementary Figure 15C). Modifying Δβ or the number of CpGs underlying the regions did not have a drastic effect on the overlap with the “truth” sets, although for the gene expression array terms, the additional filter of |Δβ| = 0.2 and number of CpGs = 4 reduced the number of “truth” gene sets identified by GOregion.

## Discussion

Gene set testing is a useful tool to gain additional biological insight into the underlying mechanisms in an experiment. Here we present GOmeth for performing gene set testing after a probe-level analysis, and GOregion, a gene set testing method following a region-based analysis. To the best of our knowledge, GOregion is the only method that specifically tests enrichment of gene sets for differentially methylated regions.

Through our investigation of the composition of the 450K and EPIC arrays, we have observed that annotating statistically significant differentially methylated CpGs to genes is not a trivial exercise. For our gene set testing methods we choose to rely on the annotation provided by Illumina, however other studies have used alternative approaches, such as defining promoter regions as regions of + / - 2kb of the transcription start sites of genes (Pidsley et al. 2016).

We have shown that two sources of bias affect gene set testing: probe-number and multi-gene bias, and our methods account for both of these. Through the use of simulations and resampling normal samples we have shown that GOmeth correctly controls the false discovery rate with minimal bias. We applied GOmeth to sorted blood cell types and developing B-cell datasets and showed that the top ranked categories are consistently biologically relevant across multiple comparisons. We defined two different types of “truth” sets based on the information available in the GO and KEGG databases, as well as independently analysed gene expression datasets. We acknowledge that our “truth” sets will have shortcomings in that they are unlikely to encompass all the truly enriched pathways, and, particularly in the case of the gene expression data analysis, the choice of gene set testing method is likely to play a role in how gene sets are ranked. Nevertheless, we showed that GOmeth generally outperforms other available methods. Further, we have shown that the probe-number bias affects the probability that a region is called differentially methylated. We therefore developed GOregion to perform unbiased gene set testing following a region-based analysis. For the blood cell types and developing B-cell dataset, GOregion outperformed a simple hypergeometric testing approach that has been previously used in analyses.

An important consideration when using GOmeth is how many differentially methylated probes to use as input to the method. For comparisons that have tens of thousands of significantly differentially methylated CpGs, in the first instance, we would recommend performing a region-based analysis and using GOregion to perform gene set testing. Another option would be to use GOmeth, but restrict the input CpGs to the top ranked CpGs, with a rule of thumb that the number of input CpGs is less than 10,000. Simply taking a false discovery rate cut-off for the B-cells vs NK cells comparison, for example, resulted in too many genes being identified as differentially methylated (>12,000) and we found that the gene set testing results were not very specific or biologically meaningful. In our comparisons, we have used the top 5000 differentially methylated probes and this produced good results for the blood cell type dataset. There is additional functionality in GOmeth to restrict the list of significant CpGs by genomic features, such as “TSS1500”, “TSS200” and “Body”, for example. This has the effect of decreasing the overall numbers of CpGs to use as input, but potentially retains more biologically meaningful loci.

For users who are not comfortable with setting thresholds to select appropriate CpGs as input to GOmeth following a probe-level analysis, mGLM and ebGSEA are alternative statistical methods that do not require setting cut-offs to select significant CpGs. Generally, when the numbers of significant probes was large, ebGSEA ranked large, broad GO terms more highly compared to mGLM. However, when the numbers of statistically significant CpGs was fewer than 5000, ebGSEA performed well in our evaluations.

Even though a region-based approach can potentially select fewer CpGs, in our analysis of the sorted blood cell types and developing B-cell data, it always identified additional unique CpGs not detected using the probe-wise methods. This is likely because these CpGs were not statistically significant on their own but are identified as part of a region. Thus, given the potential for capturing biologically important CpGs, which may be missed by probe-wise approaches due to their reliance on rankings and significance cut-offs, we suggest that a good quality, region-based analysis can potentially distil more focused gene sets than a probe-level gene set analysis of the same data.

An important feature of GOmeth and GOregion is their flexibility. Unlike the *methylGSA* methods, GOmeth and GOregion do not require any filtering of gene sets to produce robust results. GOregion is compatible with the results of any software for finding differentially methylated regions which can be expressed as a ranged data object. Both GOmeth and GOregion are computationally efficient and can test a variety of gene sets such as GO categories, KEGG pathways or any list of custom gene sets.

### Conclusions

GOmeth and GOregion are novel statistical methods that account for biases in gene set testing for methylation arrays. We have shown that our methods produce the most biologically meaningful results while controlling the false discovery rate. All of our gene set testing functions are available in the *missMethyl* Bioconductor R package.

## Methods

All analysis code presented in this manuscript can be found at http://oshlacklab.com/methyl-geneset-testing/. The analysis website was created using the *workflowr* (1.6.2) R package (Blischak, Carbonetto, and Stephens 2019). The GitHub repository associated with the analysis website is at: https://github.com/Oshlack/methyl-geneset-testing.

### Statistical model for GOmeth

The statistical test for GOmeth and GOregion is based on Wallenius’ noncentral hypergeometric distribution, which is a generalised version of the hypergeometric distribution where items are sampled with bias. For GOmeth, we take the following stepwise procedure:

1. For each CpG *i* annotated to gene *j*, calculate a weight

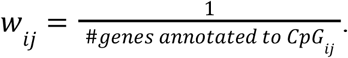

If each CpG is annotated to exactly one gene, then *w*_*ij*_ = 1.
2. Let *i = 1*, …,*I*_j_ denote the CpGs annotated to gene *j*. Calculate the equivalent number of CpGs measured across gene *j* as:

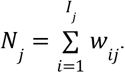 If there are no multi-gene associated CpGs, then *N*_*j*_ is simply the number of probes measured across gene *j*.
3. Let *A* define the set of significant differentially methylated Cpgs. For each gene *j*, define an indicator vector **1**_*j*_ (*x*) of length *I*_*j*_ such that *x*_*i*_ = 1 if *CpG*_*ij*_ ∈ *A*, and *x*_*i*_ = 0 if *CpG*_*ij*_ ∉ *A*, where *i = 1*, …,*I*_*j*_.
4. Let ***w***_*j*_ define the vector of weights *w*_*ij*_ for each gene *j*. Calculate the differential methylation score for each gene *j*

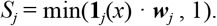 Note the maximum value of *S*_*j*_ is 1, and *S*_*j*_ < 1 only in cases where the significant CpGs are all multi-gene associated, and the summed weights are less than one. *S*_*j*_ = 0 when there are no significant differentially methylated CpGs across gene *j*.
5. Let *j* = 1, …, *J*_*g*_ denote the genes that are present in gene set *g*. Calculate the enrichment statistic for Wallenius’ noncentral hypergeometric test for each gene set *g*

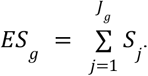 For the standard Wallenius’ noncentral hypergeometric test the enrichment statistic is the intersection between the significant differentially methylated genes and genes in gene set *g*, which ignores multi-gene associated CpGs. Our modified enrichment statistic *ES*_*g*_ accounts for multi-gene associated CpGs for gene set *g*.
6. Calculate the probability weighting function (PWF) by applying a moving average smoother to an ordered binary vector (based on the number of associated CpGs) where 1 indicates a gene is differentially methylated and 0 indicates the gene is not differentially methylated. We use the ‘tricubeMovingAverage’ function in the *limma* package which is similar to a least squares loess curve of degree zero. The binary vector is ordered by the number of equivalent CpGs measuring methylation across each gene, *N*_*j*_, from smallest to largest. The output is a vector of the same length as the input such that each gene is assigned a probability of differential methylation based on the smoothed value. We then calculate the expected odds of enrichment for each gene set *g* by calculating the mean PWF of the genes in the set and comparing it to the mean PWF of the rest of the genes represented on the array.

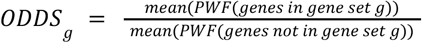
7. For testing enrichment of each gene set *g*, we obtain a one-sided p-value from Wallenius’ noncentral hypergeometric distribution with the following parameters: *x* = ⌊*ES*_*g*_⌋, *m*_*1*_ = the size of the gene set *J*_*g*_, *m*_*2*_ = the number of genes on the rest of the array, *n* = the total number of significant genes, and *odds* = *ODDS*_*g*_. We use the *BiasedUrn* R package to obtain p-values.

### Null simulations: random sampling of CpGs

We randomly selected sets of 50, 100, 500, 1000, 5000 and 10,000 CpGs from the Illumina array annotation for both 450k and EPIC arrays. The sampling was repeated 100 times for each CpG set size. We tested for significant enrichment of GO categories using a standard hypergeometric test (HGT), a Wallenius’ hypergeometric test accounting for probe number bias (HGT-mod) and GOmeth, which is based on Wallenius’ hypergeometric test and accounts for probe number and multi-gene bias.

### Methylation Datasets

The normal samples from the KIRC TCGA dataset (Cancer Genome Atlas Research Network 2013) were used for estimating the false discovery rate of the different gene set testing methods. The data was downloaded using the *curatedTCGAData* Bioconductor package (Ramos 2020) and the 160 normal samples extracted. The data was provided as already processed βvalues, however we performed additional filtering and removed poor quality probes, probes containing SNPs as well as sex chromosome probes. The resulting multidimenational scaling plots showed no apparent evidence of sex or other technical effects (Figure 3C).

In addition, 85 normal samples from the BRCA TCGA dataset were used to estimate the false discovery rates of the seven gene set testing methods. The data was downloaded using the *curatedTCGAData* Bioconductor package and the 97 normal samples extracted. Following quality control 12 samples were removed (8 with unusual beta value distributions and 4 African/African American samples). Poor quality probes and probes containing SNPs were filtered out. Probes located on the sex chromosomes were retained as all the samples were female (Supplementary Figure 3C).

To compare the performance between different gene set testing methods when there is significant differential methylation, we used Illumina Infinium HumanMethylationEPIC (GSE110554) data generated from flow-sorted neutrophils (Neu, n = 6), monocytes (Mono, n = 6), B-lymphocytes (Bcells, n = 6), CD4+ T-cells (CD4T, n=7, six samples and one technical replicate), CD8+ T-cells (CD8T, n = 6), Natural Killer cells (NK, n = 6) and 12 DNA artificial mixtures (labeled as MIX) (Salas et al. 2018). Only the sorted cells were used in our analysis. The data was downloaded using the *ExperimentHub* Bioconductor package.

We also analysed a developing B-cell dataset that had matched DNA methylation and gene expression measurements (Lee et al. 2012). Four populations of early B-cell developmental stages were obtained from human fetal bone marrow, from 8 individuals. Methylation was measured using the Illumina HumanMethylation450 Beadchip and gene expression was measured using the GeneChip Human Gene 1.0 ST Array (Affymetrix). The four populations were identified using flow cytometry antibodies and consist of Stage 1 (predominantly multipotent progenitors and common lymphoid progenitors), Stage 2 (pre-B-I cells), Stage 3 (pre-B-II cells) and Stage 4 (immature B-cells). The data was downloaded from the Gene Expression Omnibus (GSE45461).

### Analysis and processing

The majority of the analysis was performed using R (4.0.3) and some using R (3.6.1) (R Core Team 2014). The specific R and package versions used for different aspects of the analysis can be viewed under the “Session Information” sections of the analysis website associated with this study: http://oshlacklab.com/methyl-geneset-testing/.

### Quality control and normalization

All methylation data was processed using the *minfi* (Aryee et al. 2014; Fortin, Triche, and Hansen 2017) R Bioconductor (R. C. Gentleman et al. 2004; Huber et al. 2015) package. Between array and probe-type normalization was performed using the stratified quantile normalisation (SQN) method (Touleimat and Tost 2012). Probes with a detection P-value > 0.01 in one or more samples were discarded. Probes potentially affected by common SNPs (minor allele frequency > 0) proximal to the CpG of interest (up to 2 bp upstream and 1 downstream) and non-specific probes (Pidsley et al. 2016; Y.-A. Chen et al. 2013) were also removed from further analysis.

### Statistical analysis

The proportion of methylation at each CpG is represented by the βvalue, defined as the proportion of the methylated signal to the total signal and calculated from the normalized intensity values. Statistical analyses were performed on M-values 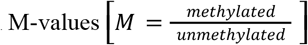 as recommended by Du et al. (2010).

### Comparison of gene set testing methods using sorted blood cell data

CpG probe-wise linear models were fitted to determine differences in methylation between cell types (B-cells vs NK, CD4 vs CD8 T-cells, monocytes vs neutrophils) using the *limma* package (Ritchie et al. 2015). Differentially methylated probes (DMPs) were identified using empirical Bayes moderated t-tests (Smyth 2005), performing robust empirical Bayes shrinkage of the gene-wise variances to protect against hypervariable probes (Phipson et al. 2016). Empirical Bayes moderated-t p-values were then calculated relative to a minimum meaningful log-fold-change (lfc) threshold on the M-value scale (lfc = 0.5, corresponding to |Δβ| ∼ 0.1) (McCarthy and Smyth 2009). P-values were adjusted for multiple testing using the Benjamini-Hochberg procedure (Benjamini and Hochberg 1995).

For each comparison, we tested for significant enrichment of GO categories and KEGG pathways. We only tested sets with at least 5 genes and at most 5000 genes for the *methylGSA* methods. The top ranked 5000 CpGs were tested using the HGT and GOmeth, from the *missMethyl* package, for enrichment of GO terms and KEGG pathways. The raw p-values were passed as input to the *methylGSA* methods. The ebGSEA methods were run using the *ebGSEA* R package (https://github.com/aet21/ebGSEA).

### Comparison of gene set testing methods using kidney clear cell carcinoma (KIRC) data

The KIRC data from the *curatedTCGAData* package was provided as β values with masked data points; data points were masked as “NA” if their detection p-value was greater than 0.05 or the probe was annotated as having a SNP within 10 base pairs or repeat within 15 base pairs of the interrogated CpG (Cancer Genome Atlas Research Network et al. 2016). We extracted only the 160 normal samples and removed probes with any “NA” values, as well as SNP-affected probes and multi-mapping and sex-chromosome probes, as previously described. This left 364,602 probes for downstream analysis.

We ran 100 null simulations by randomly subsampling the normal samples and splitting them into two artificial “groups” with 5, 10, 20, 40 and 80 samples per “group”. For each of the 100 simulations, at each sample size, DMPs between “groups” were identified using empirical Bayes moderated t-tests (Smyth 2005), performing robust empirical Bayes shrinkage of the gene-wise variances to protect against hypervariable probes (Phipson et al. 2016).

We then performed gene set testing of the differential methylation analysis results using several methods with the Broad MSigDB gene sets available in the *ChAMP* Bioconductor package. GOmeth was run using both the top 1000 and top 5000 significant CpGs as input. The *methylGSA* methods; mGLM, mRRA (ORA) and mRRA (GSEA) were run with gene set sizes restricted to a minimum of 5 and maximum of 5000 genes. The ebGSEA method was run using default parameters and both its KPMT and WT output were compared.

### Comparison of gene set testing methods using breast invasive carcinoma (BRCA) data

As with the KIRC data, the BRCA data was downloaded using the *curatedTCGAData* package. We then extracted the 97 normal samples and removed probes with any “NA” values, as well as SNP-affected probes and multi-mapping probes, as previously described. Sex chromosome probes were retained since all sample donors were female. This left 371,389 probes for further analysis. Twelve outlying samples were removed (8 with unusual beta value distributions and 4 African/African American samples), leaving 85 samples for downstream analysis.

We ran 100 null simulations by randomly subsampling the normal samples and splitting them into two artificial “groups” with 5, 10, 20, and 40 samples per “group”. DMPs between the two “groups” were identified as described for the KIRC data, followed by the same gene set testing approach using the seven different methods previously outlined.

### RNA-Seq data and analysis

The RNA-Seq data for the sorted blood cell types was downloaded from SRA (GSE107011; SRP125125) (Monaco et al. 2019; W. Xu et al. 2019). The reads were mapped to hg19 reference transcriptome (http://refgenomes.databio.org/v2/asset/hg19_cdna/fasta/archive?tag=default) and quantified using Salmon (1.2.1) (Patro et al. 2017). Salmon transcript-level estimates were imported and summarised at the gene-level as length-scaled TPM using the *tximport* Bioconductor package (Soneson, Love, and Robinson 2015). Lowly expressed genes were filtered out using the *edgeR* (Robinson, McCarthy, and Smyth 2010) *‘*filterByExpr’ function as described by Chen at al. (2016). The data was then TMM normalised (Robinson and Oshlack 2010) and transformed using ‘voomWithQualityWeights’ (Liu et al. 2015), to increase power by combining ‘voom’ (Law et al. 2014) observational-level weights with sample-specific weights.

Probe-wise linear models were then fitted for each gene to determine gene expression differences between cell types (B-cells vs NK cells, CD4 vs CD8 T-cells, monocytes vs neutrophils) using *limma* (Ritchie et al. 2015). Differentially expressed genes were identified using empirical Bayes moderated t-tests (Smyth 2005), performing robust empirical Bayes shrinkage of the gene-wise variances to protect against hypervariable probes (Phipson et al. 2016). P-values were adjusted for multiple testing using the Benjamini-Hochberg procedure (Benjamini and Hochberg 1995).

We used the ‘goana’ function from the *limma* package to test enrichment of GO categories, ‘kegga’ to test for enrichment of KEGG pathways and a generalised version of ‘goana’ and ‘kegga’; to test for enrichment of the Broad MSigDB gene sets. All the methods took gene length bias into account (Young et al. 2010). GO, KEGG and MSigDB “truth” sets were then defined for each cell type comparison from the RNA-Seq analysis as the top 100 enriched sets.

### Affymetrix array gene expression data and analysis

The Affymetrix Human Gene 1.0 ST Array gene expression data for pre B-cell development was downloaded from GEO (GSE45460) (Lee et al. 2012).

The raw CEL files were loaded and processed using the *oligo* Bioconductor package. The data was background corrected, normalised and summarised using Robust Multichip Average (RMA) pre-processing (Bolstad et al. 2003; Irizarry et al. 2003). Only probes with a median intensity greater than 4.5, in at least 7 samples, were retained. Transcript-cluster identifiers that mapped to multiple Entrez identifiers were filtered out, along with any probes that did not map to Entrez identifiers, leaving 19,494 genes for downstream analysis.

Probe-wise linear models were then fitted for each gene to determine gene expression differences between Stage 1 and Stage 2 of pre B-cell development (Stage 1 vs Stage 2) using *limma* (Ritchie et al. 2015). Differentially expressed genes were identified using empirical Bayes moderated t-tests (Smyth 2005), performing robust empirical Bayes shrinkage of the gene-wise variances to protect against hypervariable probes (Phipson et al. 2016). P-values were adjusted for multiple testing using the Benjamini-Hochberg procedure (Benjamini and Hochberg 1995). Genes with a log_2_ fold change greater than 0.5 using TREAT (McCarthy and Smyth 2009) and FDR less than 0.05 were deemed to be differentially expressed.

The ‘goana’ function from the *limma* package was used to test enrichment of GO categories and ‘kegga’ to test for enrichment of KEGG pathways. GO and KEGG “truth” sets were then defined for the Stage 1 versus Stage 2 comparison from the gene expression analysis as the top 100 enriched sets.

### Evaluation of GOregion using flow sorted blood cell data

The lllumina Infinium HumanMethylationEPIC (GSE110554) data generated from flow-sorted blood cells was used for identification of DMRs. The data was processed as previously described. DMRs between cell types (B-cells vs NK cells, CD4 vs CD8 T-cells, monocytes vs neutrophils) were identified using the *DMRcate* Bioconductor package (Peters et al. 2015). The analysis was performed on M-values using default parameters. Downstream gene set testing was performed on a filtered list of DMRs with a mean |Δβ| ≥ 0.1 and at least 3 underlying CpGs.

GO terms were tested for enrichment of DMR-associated genes using ‘goregion’ and a standard HGT, as implemented in the ‘goana’ function from the *limma* Bioconductor package. A gene, as defined in the *TxDb*.*Hsapiens*.*UCSC*.*hg19*.*knownGene* Bioconductor package, was included in the list of genes to be tested using ‘goana’ if it overlapped a DMR by at least 1bp.

### Evaluation of GOregion using pre B-cell development data

The lllumina Infinium HumanMethylation450 (GSE45459) data generated from 4 stages of pre B-cell development was used for identification of DMRs. The data was processed as previously described. DMRs between Stage 1 and Stage 2 were identified using the *DMRcate* Bioconductor package (Peters et al. 2015). The analysis was performed on M-values using default parameters. DMRs were not filtered prior to gene set testing.

GO terms were tested for enrichment of DMR-associated genes using ‘goregion’ and a standard HGT, as previously described for the sorted blood cell data.

## Abbreviations

TCGA: The Cancer Genome Atlas
MSigDB: Molecular Signatures Database
RNA-Seq: RNA sequencing
FDR: false discovery rate
DMP: differentially methylated probe
DMR: differentially methylated region

## Declarations

### Ethics approval and consent to participate

Not applicable.

### Consent for publication

Not applicable.

### Availability of data and materials

All data used in this manuscript is publicly available. The KIRC and BRCA data was downloaded using the *curatedTCGAData* R Bioconductor package. The sorted blood cell type Illumina Infinium HumanMethylationEPIC data is available from the Gene Expression Omnibus under accession number GSE110554 and was downloaded using the *ExperimentHub* R Bioconductor package. The RNA-Seq data for the sorted blood cell types was downloaded from SRA (GSE107011; SRP125125). The methylation and gene expression datasets for the sorted B-cell development data was downloaded from the Gene Expression Omnibus under accession number GSE45461. All analysis code presented in this manuscript can be found at the following *workflowr* website http://oshlacklab.com/methyl-geneset-testing/. The GitHub repository associated with the analysis website is at: https://github.com/Oshlack/methyl-geneset-testing.

### Competing interests

The authors declare that they have no competing interests.

### Funding

BP is supported by an Emerging Leader Investigator Grant (GNT1175653) from the National Health and Medical Research Council (NHMRC). AO is supported by NHMRC GNT1126157. The funding body did not play any role in the study design, analysis, interpretation of data or writing of the manuscript.

### Authors’ contributions

The original idea for gene set testing for methylation array data was conceived by AO. BP conceived the statistical model and wrote the core of the ‘gometh’ and ‘gsameth’ code. JM conceived the idea of extending GOmeth to DMRs. BP and JM determined how to apply GOmeth to DMRs. JM wrote the GOregion code. JM performed all analysis and produced the figures and analysis website. BP and JM wrote the initial draft of the manuscript, which was revised by all authors. All authors read and approved the final manuscript.

## Acknowledgements

We would like to acknowledge Peter Langfelder for providing detailed code and simulations which helped us to discover and correct for multi-gene bias in our testing framework. We would also like to acknowledge all *missMethyl* users who have provided valuable bug reports and feedback.

## Notes

http://oshlacklab.com/methyl-geneset-testing/

https://www.bioconductor.org/packages/release/bioc/html/missMethyl.html

### Competing Interest Statement

The authors have declared no competing interest.

### Summary of Updates

In the revised version of our manuscript, we have added the analysis of two new methylation datasets, including a dataset that has matched gene expression data. We have repeated our analyses using the new implementation of the ebGSEA methods and have incorporated these into our results and modified the figures accordingly. In addition, we have added text to clarify how the statistical framework in GOmeth is implemented.

