## Supplementary Figures for "Gene set enrichment analysis for genome-wide DNA methylation data"

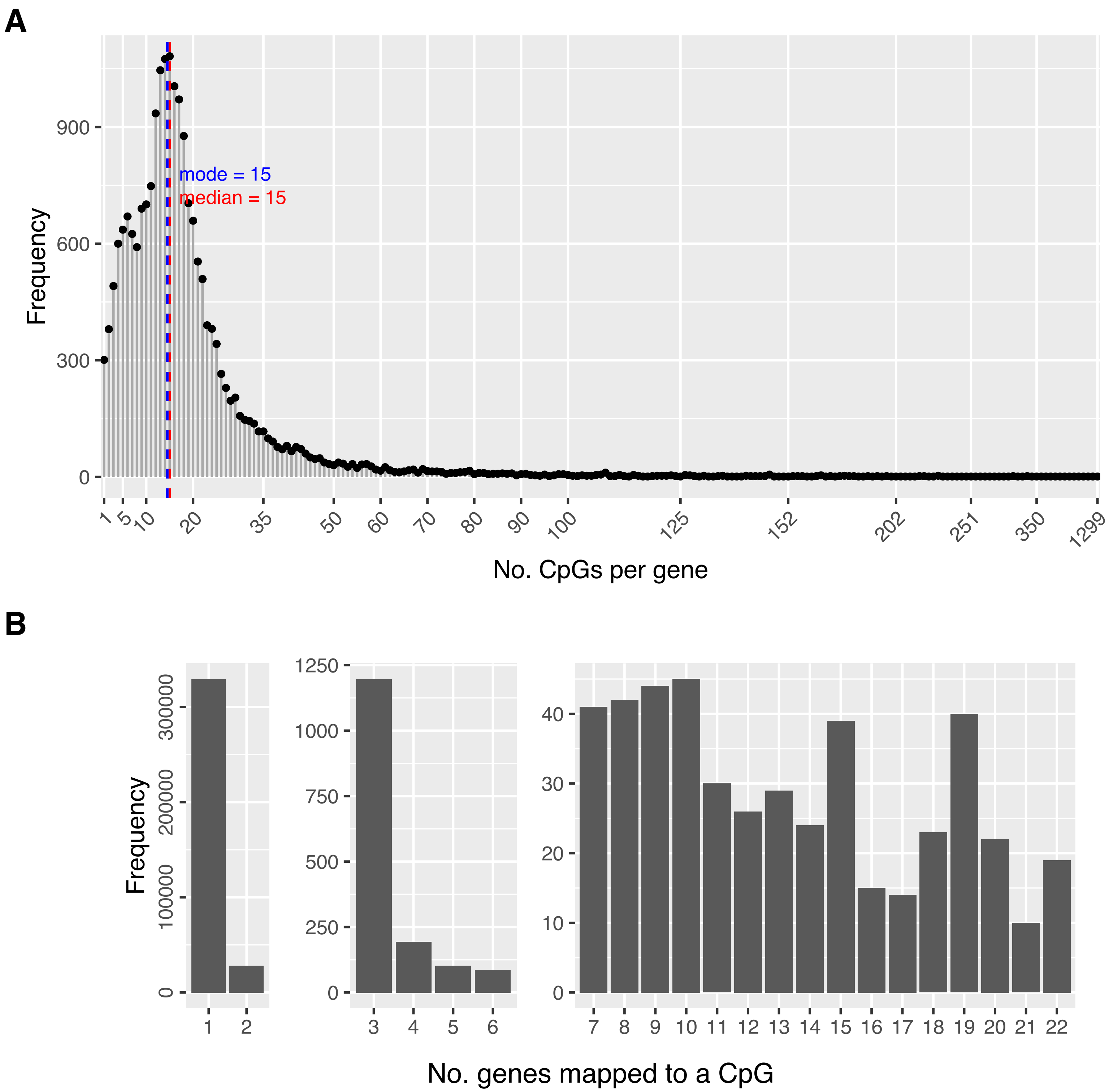
A
Frequency
900
600
300
0
mode = 15
median = 15
1
5
10
20
35
50
60
70
80
90
100
125
152
202
251
350
1299
No. CpGs per gene
B
Frequency
300000
200000
100000
0
1
2
1250
1000
750
500
250
0
3
4
5
6
40
30
20
10
0
No. genes mapped to a CpG

**Supplementary Figure 1**. **Array design bias for the Illumina HumanMethylation 450K BeadChip. (A)** Frequency plot of the numbers of CpGs measuring methylation across each gene for the 450K array (probe number bias). The most extreme value is 1299 CpGs measuring methylation for a single gene. The median is 15 and the mode is 15. **(B)** Split bar chart showing the numbers of genes annotated to each CpG (multi-gene bias). While the majority of CpGs are annotated to only one gene, there is still a large number annotated to 2 or more genes.

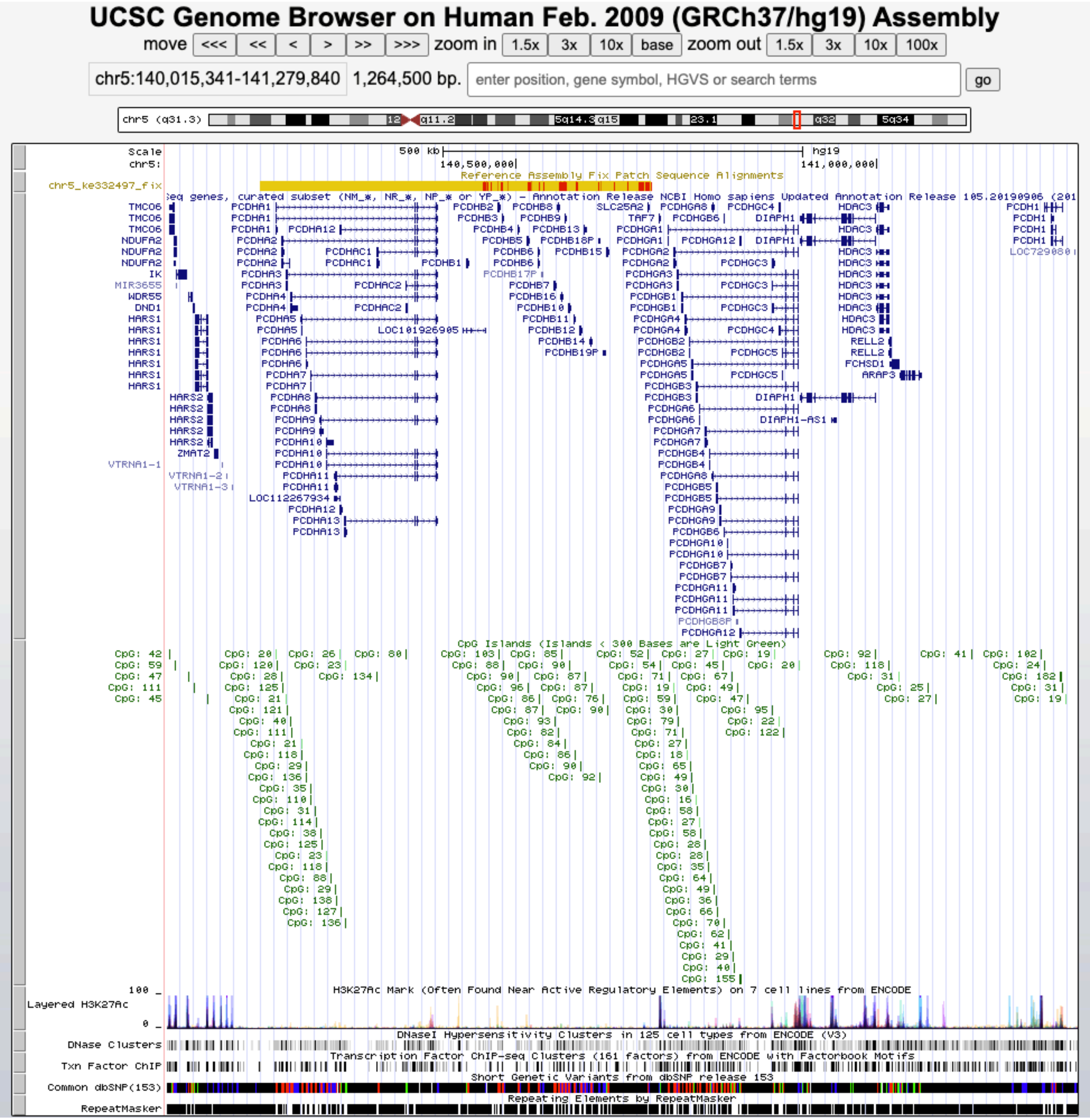
UCSC Genome Browser on Human Feb. 2009 (GRCh37/hg19) Assembly
move <<< << < > >> >>> zoom in 1.5x 3x 10x base zoom out 1.5x 3x 10x 100x
chr5:140,015,341-141,279,840 1,264,500 bp.
enter position, gene symbol, HGVS or search terms
go
chr5 (q31.3)
Scale
chr5:
500 kb
hg19
140,500,000
141,000,000
Reference Assembly Fix Patch Sequence Alignments
chr5_ke332497_fix
CpG Islands (Islands < 300 Bases are Light Green)
H3K27Ac Mark (Often Found Near Active Regulatory Elements) on 7 cell lines from ENCODE
Layered H3K27Ac
DNaseI Hypersensitivity Clusters in 125 cell types from ENCODE (V3)
DNase Clusters
Transcription Factor ChIP-seq Clusters (161 factors) from ENCODE with Factorbook Motifs
Txn Factor ChIP
Short Genetic Variants from dbSNP release 153
Common dbSNP(153)
Repeating Elements by RepeatMasker
RepeatMasker

**Supplementary Figure 2.** UCSC genome browser snapshot of the protocadherin gamma cluster.

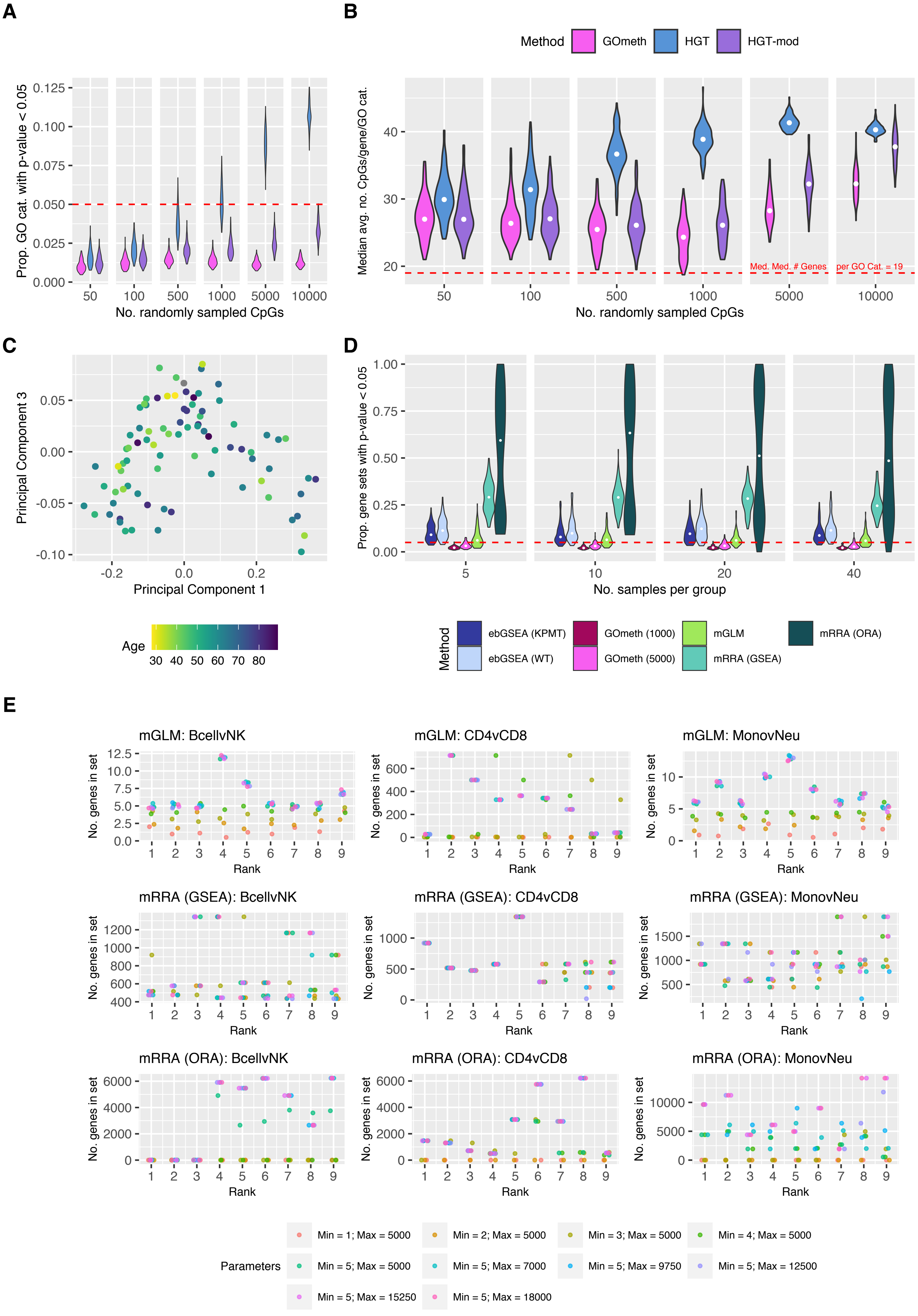
A
B
C
D
E
Method
GOmeth
HGT
HGT-mod
Prop. GO cat. with p-value < 0.05
No. randomly sampled CpGs
Median avg. no. CpGs/gene/GO cat.
Med. Med. # Genes per GO Cat. = 19
Principal Component 3
Principal Component 1
Age
Prop. gene sets with p-value < 0.05
No. samples per group
ebGSEA (KPMT)
ebGSEA (WT)
GOmeth (1000)
GOmeth (5000)
mGLM
mRRA (GSEA)
mRRA (ORA)
mGLM: BcellvNK
mGLM: CD4vCD8
mGLM: MonovNeu
mRRA (GSEA): BcellvNK
mRRA (GSEA): CD4vCD8
mRRA (GSEA): MonovNeu
mRRA (ORA): BcellvNK
mRRA (ORA): CD4vCD8
mRRA (ORA): MonovNeu
No. genes in set
Rank
Parameters
Min = 1; Max = 5000
Min = 2; Max = 5000
Min = 3; Max = 5000
Min = 4; Max = 5000
Min = 5; Max = 5000
Min = 5; Max = 7000
Min = 5; Max = 9750
Min = 5; Max = 12500
Min = 5; Max = 15250
Min = 5; Max = 18000

**Supplementary Figure 3. Evaluation of false discovery rate control for 450K data. (A)** Type I error rates across 100 simulations for varying numbers of randomly sampled CpGs. **(B)** Median average numbers of CpGs per gene for GO categories with an unadjusted p-value < 0.05. The hypergeometric test is biased towards GO categories with more CpGs per gene on average. GOmeth = adjust for probe-number and multi-gene bias; HGT = hypergeometric test; HGT-mod = adjust for probe-number bias only. **(C)** Multidimensional scaling plot of normal samples from TCGA BRCA data, coloured by age. **(D)** False discovery rate control of seven gene set testing methods using normal samples from TCGA BRCA data. Two groups were generated by randomly sampling *n* samples per group, followed by differential methylation analysis and subsequent gene set testing. This was repeated 100 times at each sample size. The proportion of gene sets with unadjusted p-value < 0.05 across the 100 null simulations is shown for each method, at each sample size. Methods with good false discovery rate control should have relatively tight distributions around the red dashed line at 0.05. ebGSEA (KPMT) = ebGSEA using Known Population Median Test; ebGSEA (WT) = ebGSEA using Wilcoxon Test; GOmeth (1000) = GOmeth using top 1000 ranked probes; GOmeth (5000) = GOmeth using top 5000 ranked probes; mGLM = methylglm; mRRA (GSEA) = methylRRA using gene set enrichment analysis; mRRA (ORA) = methylRRA using over-representation analysis.**(E)** Gene set testing was performed on the results of the three blood cell type comparisons: CD4 T-cells vs. CD8 T-cells, monocytes vs. neutrophils and B-cells vs. NK cells, using the *MethylGSA* methods: mGLM, mRRA (GSEA) and mRRA (ORA), with several combinations of minimum and maximum gene set size parameters. When the minimum gene set size is set to less than five, mGLM and mRRA (ORA) highly rank very small gene sets containing the number of genes equal to the minimum size parameter. mRRA is also biased towards highly ranking very large gene sets, if they are not filtered out. mGLM = methylglm; mRRA (GSEA) = methylRRA using gene set enrichment analysis; mRRA (ORA) = methylRRA using over-representation analysis.

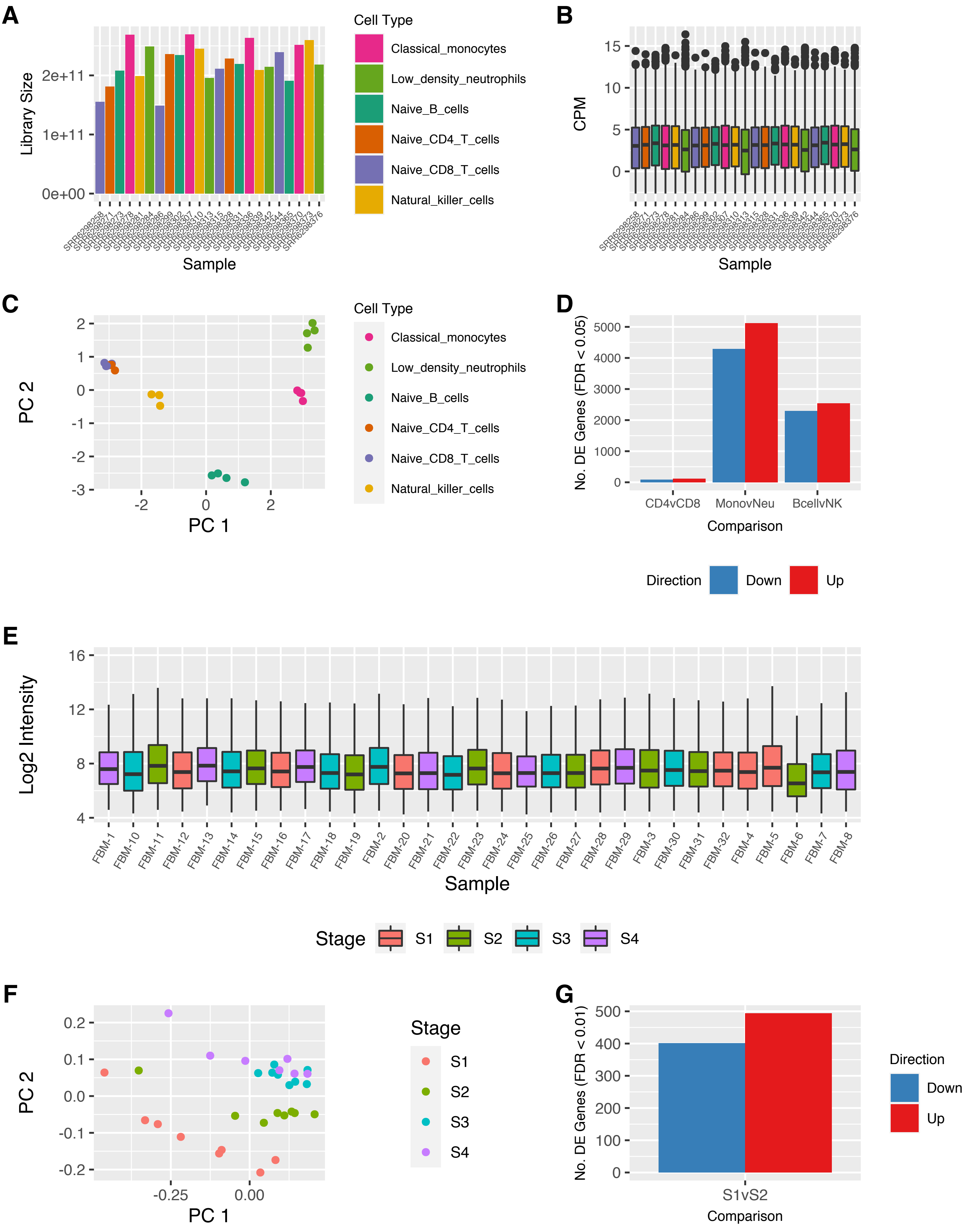
A
Library Size
2e+11
1e+11
0e+00
Sample
Cell Type
Classical_monocytes
Low_density_neutrophils
Naive_B_cells
Naive_CD4_T_cells
Naive_CD8_T_cells
Natural_killer_cells
B
CPM
15
10
5
0
Sample
C
PC 2
PC 1
Cell Type
Classical_monocytes
Low_density_neutrophils
Naive_B_cells
Naive_CD4_T_cells
Naive_CD8_T_cells
Natural_killer_cells
D
No. DE Genes (FDR < 0.05)
5000
4000
3000
2000
1000
0
CD4vCD8
MonovNeu
BcellvNK
Comparison
Direction
Down
Up
E
Log2 Intensity
16
12
8
4
FBM-1
FBM-10
FBM-11
FBM-12
FBM-13
FBM-14
FBM-15
FBM-16
FBM-17
FBM-18
FBM-19
FBM-2
FBM-20
FBM-21
FBM-22
FBM-23
FBM-24
FBM-25
FBM-26
FBM-27
FBM-28
FBM-29
FBM-3
FBM-30
FBM-31
FBM-32
FBM-4
FBM-5
FBM-6
FBM-7
FBM-8
Sample
Stage
S1
S2
S3
S4
F
PC 2
0.2
0.1
0.0
-0.1
-0.2
-0.25
0.00
PC 1
Stage
S1
S2
S3
S4
G
No. DE Genes (FDR < 0.01)
500
400
300
200
100
0
S1vS2
Comparison
Direction
Down
Up

**Supplementary Figure 4. Characteristics of gene expression datasets and analysis. (A)** Flow sorted blood cell types RNA-seq data: Library sizes (length-scaled TPM) for each of the samples after Salmon mapping and quantification. **(B)** Flow sorted blood cell types RNA-seq data: Distribution of counts per million (CPM) after filtering and normalisation. **(C)** Flow sorted blood cell types RNA-seq data: Multidimensional scaling plot of the filtered and normalised RNAseq data. **(D)** Flow sorted blood cell types RNA-seq data: Numbers of differentially expressed genes with an adjusted p-value < 0.05, for each cell type comparison: CD4 T-cells vs. CD8 T-cells, monocytes vs. neutrophils and B-cells vs. NK cells. The blue bar is the number of significantly down-regulated genes, and the red bar is the number that are significantly up-regulated; e.g. ~2,300 genes are down-regulated and ~2500 are up-regulated in B-cells, compared to NK cells. (E) Log2 intensity distributions of the raw data for all B-cell development samples. (F) Multidimensional scaling plot of the normalised and filtered B-cell development Affymetrix array data. (G) Numbers of differentially expressed genes with an adjusted p-value < 0.01, for Stage 1 versus Stage 2 of B-cell development. The blue bar is the number of significantly down-regulated genes, and the red bar is the number that are significantly up-regulated; e.g. ~400 genes are down-regulated and ~500 are up-regulated in Stage 1, compared to Stage 2.

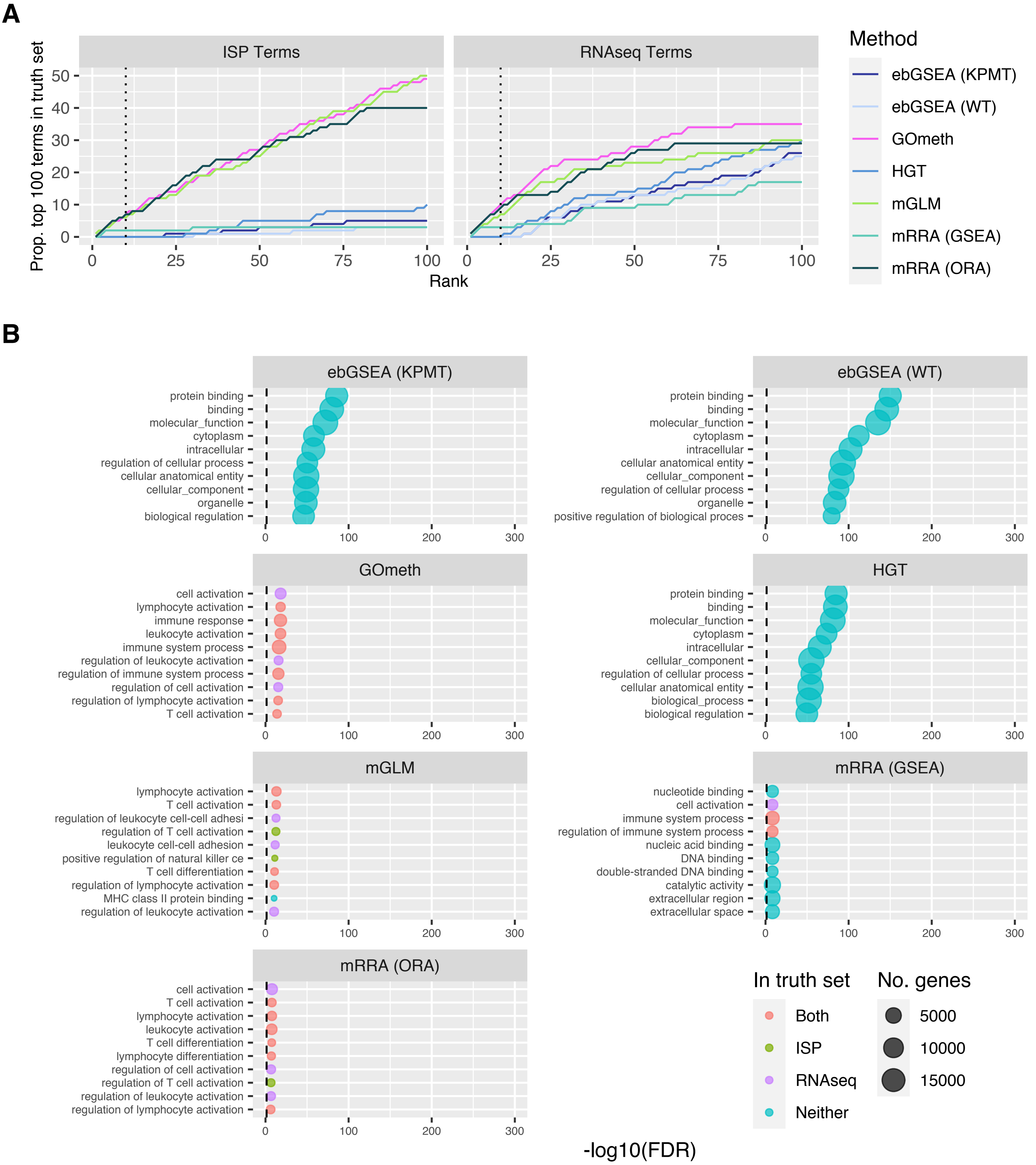

A
ISP Terms
RNAseq Terms
Prop. top 100 terms in truth set
Rank
Method
ebGSEA (KPMT)
ebGSEA (WT)
GOmeth
HGT
mGLM
mRRA (GSEA)
mRRA (ORA)
B
ebGSEA (KPMT)
protein binding
binding
molecular_function
cytoplasm
intracellular
regulation of cellular process
cellular anatomical entity
cellular_component
organelle
biological regulation
ebGSEA (WT)
protein binding
binding
molecular_function
cytoplasm
intracellular
cellular anatomical entity
cellular_component
regulation of cellular process
organelle
positive regulation of biological proces
GOmeth
cell activation
lymphocyte activation
immune response
leukocyte activation
immune system process
regulation of leukocyte activation
regulation of immune system process
regulation of cell activation
regulation of lymphocyte activation
T cell activation
HGT
protein binding
binding
molecular_function
cytoplasm
intracellular
cellular_component
regulation of cellular process
cellular anatomical entity
biological_process
biological regulation
mGLM
lymphocyte activation
T cell activation
regulation of leukocyte cell-cell adhesi
regulation of T cell activation
leukocyte cell-cell adhesion
positive regulation of natural killer ce
T cell differentiation
regulation of lymphocyte activation
MHC class II protein binding
regulation of leukocyte activation
mRRA (GSEA)
nucleotide binding
cell activation
immune system process
regulation of immune system process
nucleic acid binding
DNA binding
double-stranded DNA binding
catalytic activity
extracellular region
extracellular space
mRRA (ORA)
cell activation
T cell activation
lymphocyte activation
leukocyte activation
T cell differentiation
lymphocyte differentiation
regulation of cell activation
regulation of T cell activation
regulation of leukocyte activation
regulation of lymphocyte activation
In truth set
Both
ISP
RNAseq
Neither
No. genes
5000
10000
15000
-log10(FDR)

**Supplementary Figure 5. Comparison of gene set testing performance for CD4 T cells vs. CD8 T-cells (GO). (A)** Cumulative number of GO terms, as ranked by various methods, that are present in each truth set for the CD4 T-cells vs. CD8 T-cells comparison. ISP Terms = immune-system process child terms truth set; RNAseq Terms = top 100 terms from RNAseq analysis of the same cell types. **(B)** Bubble plots of the top 10 GO terms as ranked by various gene set testing methods for the CD4 T-cells vs. CD8 T-cells comparison. The size of the bubble indicates the relative number of genes in the set. The colour of the bubble indicates whether the term is present in either RNAseq (purple) or ISP (green) truth sets, both (red) or neither (blue). ebGSEA (KPMT) = ebGSEA using Known Population Median Test; ebGSEA (WT) = ebGSEA using Wilcoxon Test; GOmeth = GOmeth using either FDR < 0.05 (for contrasts with <5000 significant CpGs) or top 5000 most significant probes; HGT = hypergeometric test; mGLM = methylglm; mRRA (GSEA) = methylRRA using gene set enrichment analysis; mRRA (ORA) = methylRRA using over-representation analysis.

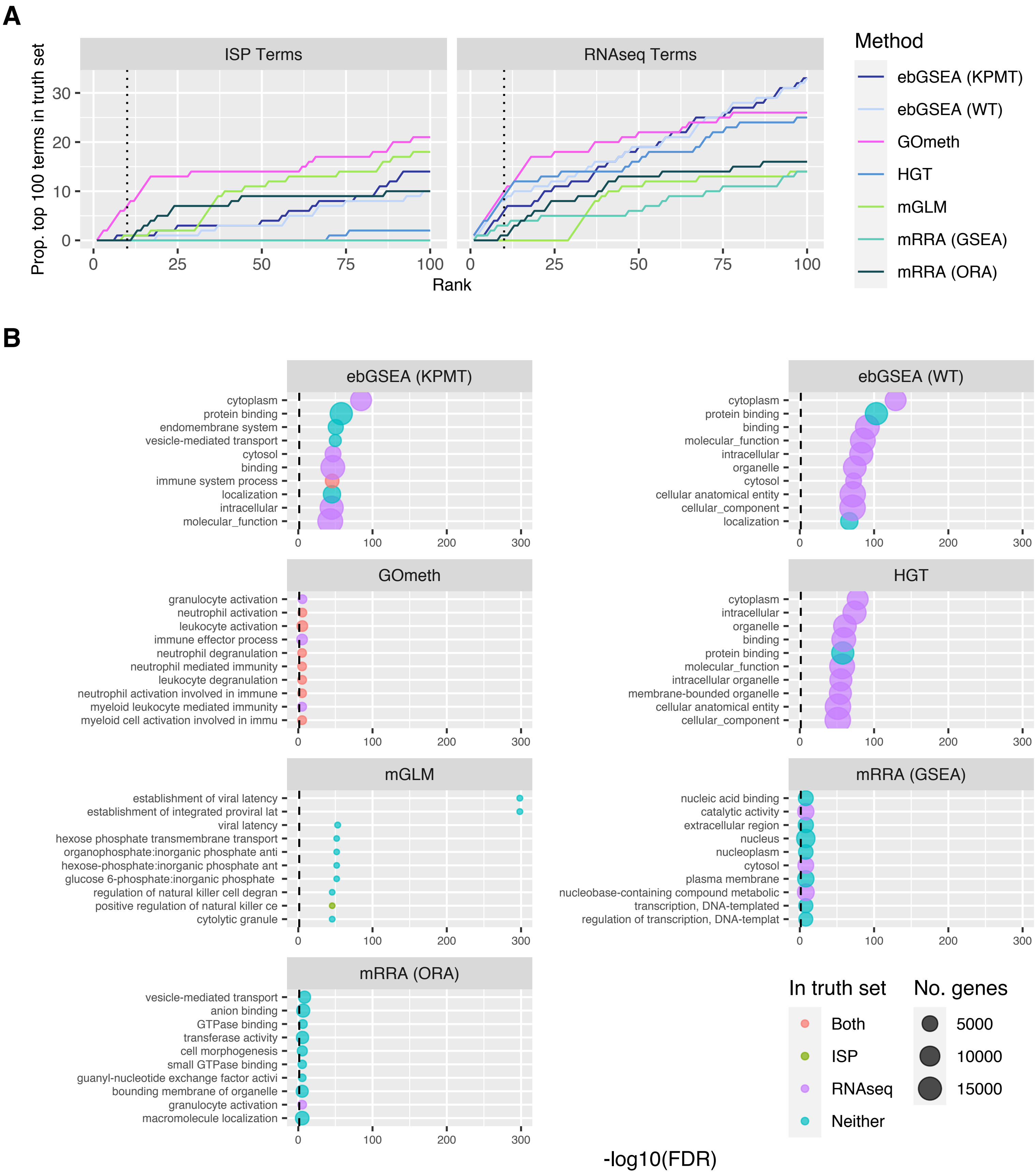
A
ISP Terms
RNAseq Terms
Prop. top 100 terms in truth set
Rank
Method
ebGSEA (KPMT)
ebGSEA (WT)
GOmeth
HGT
mGLM
mRRA (GSEA)
mRRA (ORA)
B
ebGSEA (KPMT)
cytoplasm
protein binding
endomembrane system
vesicle-mediated transport
cytosol
binding
immune system process
localization
intracellular
molecular_function
ebGSEA (WT)
cytoplasm
protein binding
binding
molecular_function
intracellular
organelle
cytosol
cellular anatomical entity
cellular_component
localization
GOmeth
granulocyte activation
neutrophil activation
leukocyte activation
immune effector process
neutrophil degranulation
neutrophil mediated immunity
leukocyte degranulation
neutrophil activation involved in immune
myeloid leukocyte mediated immunity
myeloid cell activation involved in immu
HGT
cytoplasm
intracellular
organelle
binding
protein binding
molecular_function
intracellular organelle
membrane-bounded organelle
cellular anatomical entity
cellular_component
mGLM
establishment of viral latency
establishment of integrated proviral lat
viral latency
hexose phosphate transmembrane transport
organophosphate:inorganic phosphate anti
hexose-phosphate:inorganic phosphate ant
glucose 6-phosphate:inorganic phosphate
regulation of natural killer cell degran
positive regulation of natural killer ce
cytolytic granule
mRRA (GSEA)
nucleic acid binding
catalytic activity
extracellular region
nucleus
nucleoplasm
cytosol
plasma membrane
nucleobase-containing compound metabolic
transcription, DNA-templated
regulation of transcription, DNA-templat
mRRA (ORA)
vesicle-mediated transport
anion binding
GTPase binding
transferase activity
cell morphogenesis
small GTPase binding
guanyl-nucleotide exchange factor activi
bounding membrane of organelle
granulocyte activation
macromolecule localization
In truth set
Both
ISP
RNAseq
Neither
No. genes
5000
10000
15000
-log10(FDR)

**Supplementary Figure 6. Comparison of gene set testing performance for monocytes vs. neutrophils (GO). (A)** Cumulative number of GO terms, as ranked by various methods, that are present in each truth set for the monocytes vs. neutrophils comparison. ISP Terms = immune-system process child terms truth set; RNAseq Terms = top 100 terms from RNAseq analysis of the same cell types. **(B)** Bubble plots of the top 10 GO terms as ranked by various gene set testing methods for the monocytes vs. neutrophils comparison. The size of the bubble indicates the relative number of genes in the set. The colour of the bubble indicates whether the term is present in either RNAseq (purple) or ISP (green) truth sets, both (red) or neither (blue). ebGSEA (KPMT) = ebGSEA using Known Population Median Test; ebGSEA (WT) = ebGSEA using Wilcoxon Test; GOmeth = GOmeth using either FDR < 0.05 (for contrasts with <5000 significant CpGs) or top 5000 most significant probes; HGT = hypergeometric test; mGLM = methylglm; mRRA (GSEA) = methylRRA using gene set enrichment analysis; mRRA (ORA) = methylRRA using over-representation analysis.

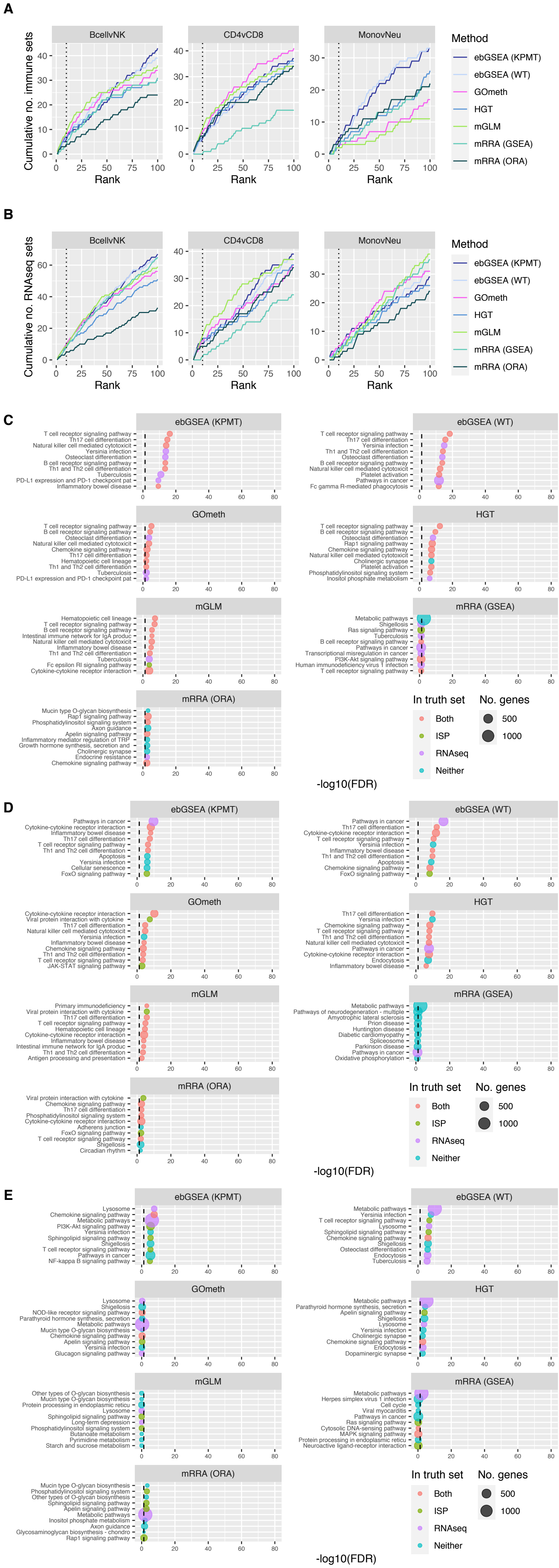

A
BcellvNK
CD4vCD8
MonovNeu
Cumulative no. immune sets
Rank
Method
ebGSEA (KPMT)
ebGSEA (WT)
GOmeth
HGT
mGLM
mRRA (GSEA)
mRRA (ORA)
B
BcellvNK
CD4vCD8
MonovNeu
Cumulative no. RNAseq sets
Rank
Method
ebGSEA (KPMT)
ebGSEA (WT)
GOmeth
HGT
mGLM
mRRA (GSEA)
mRRA (ORA)
C
ebGSEA (KPMT)
T cell receptor signaling pathway
Th17 cell differentiation
Natural killer cell mediated cytotoxicit
Yersinia infection
Osteoclast differentiation
B cell receptor signaling pathway
Th1 and Th2 cell differentiation
Tuberculosis
PD-L1 expression and PD-1 checkpoint pat
Inflammatory bowel disease
ebGSEA (WT)
T cell receptor signaling pathway
Th17 cell differentiation
Yersinia infection
Th1 and Th2 cell differentiation
Osteoclast differentiation
B cell receptor signaling pathway
Natural killer cell mediated cytotoxicit
Platelet activation
Pathways in cancer
Fc gamma R-mediated phagocytosis
GOmeth
T cell receptor signaling pathway
B cell receptor signaling pathway
Osteoclast differentiation
Natural killer cell mediated cytotoxicit
Chemokine signaling pathway
Th17 cell differentiation
Hematopoietic cell lineage
Th1 and Th2 cell differentiation
Tuberculosis
PD-L1 expression and PD-1 checkpoint pat
HGT
T cell receptor signaling pathway
B cell receptor signaling pathway
Osteoclast differentiation
Rap1 signaling pathway
Chemokine signaling pathway
Natural killer cell mediated cytotoxicit
Cholinergic synapse
Platelet activation
Phosphatidylinositol signaling system
Inositol phosphate metabolism
mGLM
Hematopoietic cell lineage
T cell receptor signaling pathway
B cell receptor signaling pathway
Intestinal immune network for IgA produc
Natural killer cell mediated cytotoxicit
Inflammatory bowel disease
Th1 and Th2 cell differentiation
Tuberculosis
Fc epsilon RI signaling pathway
Cytokine-cytokine receptor interaction
mRRA (GSEA)
Metabolic pathways
Shigellosis
Ras signaling pathway
Tuberculosis
B cell receptor signaling pathway
Pathways in cancer
Transcriptional misregulation in cancer
PI3K-Akt signaling pathway
Human immunodeficiency virus 1 infection
T cell receptor signaling pathway
mRRA (ORA)
Mucin type O-glycan biosynthesis
Rap1 signaling pathway
Phosphatidylinositol signaling system
Axon guidance
Apelin signaling pathway
Inflammatory mediator regulation of TRP
Growth hormone synthesis, secretion and
Cholinergic synapse
Endocrine resistance
Chemokine signaling pathway
In truth set
Both
ISP
RNAseq
Neither
No. genes
500
1000
-log10(FDR)
D
ebGSEA (KPMT)
Pathways in cancer
Cytokine-cytokine receptor interaction
Inflammatory bowel disease
Th17 cell differentiation
T cell receptor signaling pathway
Th1 and Th2 cell differentiation
Apoptosis
Yersinia infection
Cellular senescence
FoxO signaling pathway
ebGSEA (WT)
Pathways in cancer
Th17 cell differentiation
Cytokine-cytokine receptor interaction
T cell receptor signaling pathway
Yersinia infection
Inflammatory bowel disease
Th1 and Th2 cell differentiation
Apoptosis
Chemokine signaling pathway
FoxO signaling pathway
GOmeth
Cytokine-cytokine receptor interaction
Viral protein interaction with cytokine
Th17 cell differentiation
Natural killer cell mediated cytotoxicit
Yersinia infection
Inflammatory bowel disease
Chemokine signaling pathway
Th1 and Th2 cell differentiation
T cell receptor signaling pathway
JAK-STAT signaling pathway
HGT
Th17 cell differentiation
Yersinia infection
Chemokine signaling pathway
T cell receptor signaling pathway
Th1 and Th2 cell differentiation
Natural killer cell mediated cytotoxicit
Pathways in cancer
Cytokine-cytokine receptor interaction
Endocytosis
Inflammatory bowel disease
mGLM
Primary immunodeficiency
Viral protein interaction with cytokine
Th17 cell differentiation
T cell receptor signaling pathway
Hematopoietic cell lineage
Cytokine-cytokine receptor interaction
Inflammatory bowel disease
Intestinal immune network for IgA produc
Th1 and Th2 cell differentiation
Antigen processing and presentation
mRRA (GSEA)
Metabolic pathways
Pathways of neurodegeneration - multiple
Amyotrophic lateral sclerosis
Prion disease
Huntington disease
Diabetic cardiomyopathy
Spliceosome
Parkinson disease
Pathways in cancer
Oxidative phosphorylation
mRRA (ORA)
Viral protein interaction with cytokine
Chemokine signaling pathway
Th17 cell differentiation
Phosphatidylinositol signaling system
Cytokine-cytokine receptor interaction
Adherens junction
FoxO signaling pathway
T cell receptor signaling pathway
Shigellosis
Circadian rhythm
In truth set
Both
ISP
RNAseq
Neither
No. genes
500
1000
-log10(FDR)
E
ebGSEA (KPMT)
Lysosome
Chemokine signaling pathway
Metabolic pathways
PI3K-Akt signaling pathway
Yersinia infection
Sphingolipid signaling pathway
Shigellosis
T cell receptor signaling pathway
Pathways in cancer
NF-kappa B signaling pathway
ebGSEA (WT)
Metabolic pathways
Yersinia infection
T cell receptor signaling pathway
Lysosome
Sphingolipid signaling pathway
Chemokine signaling pathway
Shigellosis
Osteoclast differentiation
Endocytosis
Tuberculosis
GOmeth
Lysosome
Shigellosis
NOD-like receptor signaling pathway
Parathyroid hormone synthesis, secretion
Metabolic pathways
Mucin type O-glycan biosynthesis
Chemokine signaling pathway
Apelin signaling pathway
Yersinia infection
Glucagon signaling pathway
HGT
Metabolic pathways
Parathyroid hormone synthesis, secretion
Apelin signaling pathway
Shigellosis
Lysosome
Yersinia infection
Cholinergic synapse
Chemokine signaling pathway
Endocytosis
Dopaminergic synapse
mGLM
Other types of O-glycan biosynthesis
Mucin type O-glycan biosynthesis
Protein processing in endoplasmic reticu
Lysosome
Sphingolipid signaling pathway
Long-term depression
Phosphatidylinositol signaling system
Butanoate metabolism
Pyrimidine metabolism
Starch and sucrose metabolism
mRRA (GSEA)
Metabolic pathways
Herpes simplex virus 1 infection
Cell cycle
Viral myocarditis
Pathways in cancer
Ras signaling pathway
Cytosolic DNA-sensing pathway
MAPK signaling pathway
Protein processing in endoplasmic reticu
Neuroactive ligand-receptor interaction
mRRA (ORA)
Mucin type O-glycan biosynthesis
Phosphatidylinositol signaling system
Other types of O-glycan biosynthesis
Sphingolipid signaling pathway
Apelin signaling pathway
Metabolic pathways
Inositol phosphate metabolism
Axon guidance
Glycosaminoglycan biosynthesis - chondro
Rap1 signaling pathway
In truth set
Both
ISP
RNAseq
Neither
No. genes
500
1000
-log10(FDR)

**Supplementary Figure 7. Comparison of gene set testing performance for sorted blood cell types data (KEGG). (A)** Cumulative number of KEGG pathways, as ranked by various methods, that are present in the immune truth set for all comparisons. Immune sets = all pathways belonging to the following categories: Immune system, Immune disease, Signal transduction, Signaling molecules and interaction. **(B)** Cumulative number of KEGG pathways, as ranked by various methods, that are present in the RNAseq truth set for all comparisons. RNAseq Terms = top 100 KEGG pathways from RNAseq analysis of the same cell types. **(C)** Bubble plots of the top 10 KEGG pathways as ranked by various gene set testing methods for the B-cell vs. NK comparison. The size of the bubble indicates the relative number of genes in the set. The colour of the bubble indicates whether the term is present in either RNAseq (purple) or ISP (green) truth sets, both (red) or neither (blue). **(D)** Bubble plots of the top 10 KEGG pathways as ranked by various gene set testing methods for the CD4 T-cells vs. CD8 T-cells comparison. **(E)** Bubble plots of the top 10 KEGG pathways as ranked by various gene set testing methods for the monocytes vs. neutrophils comparison. ebGSEA (KPMT) = ebGSEA using Known Population Median Test; ebGSEA (WT) = ebGSEA using Wilcoxon Test; GOmeth = GOmeth using either FDR < 0.05 (for contrasts with <5000 significant CpGs) or top 5000 most significant probes; HGT = hypergeometric test; mGLM = methylglm; mRRA (GSEA) = methylRRA using gene set enrichment analysis; mRRA (ORA) = methylRRA using over-representation analysis.

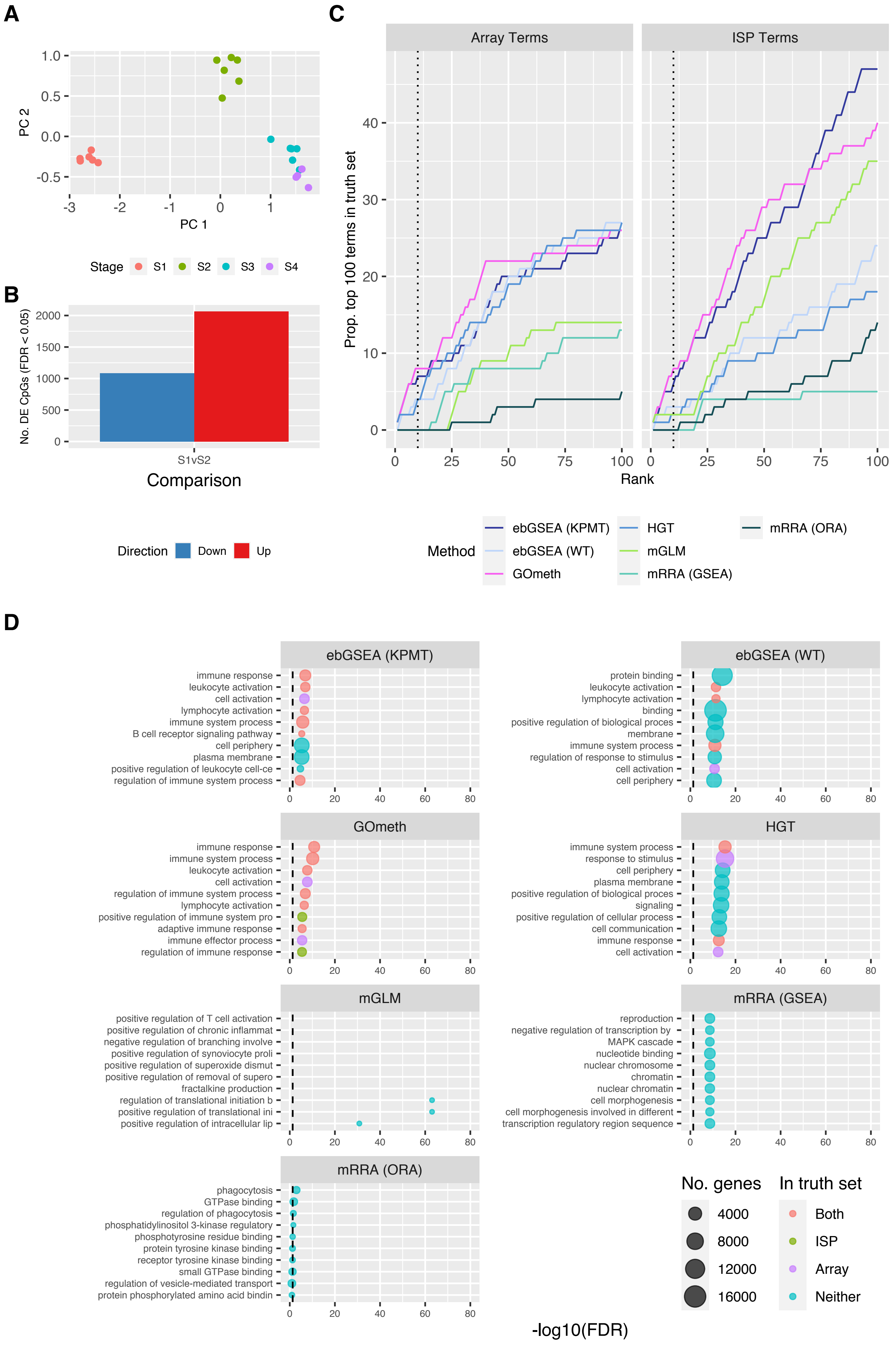
A
PC 2
PC 1
Stage S1 S2 S3 S4
B
No. DE CpGs (FDR < 0.05)
S1vS2
Comparison
Direction Down Up
C
Array Terms
ISP Terms
Prop. top 100 terms in truth set
Rank
Method
ebGSEA (KPMT)
ebGSEA (WT)
GOmeth
HGT
mGLM
mRRA (GSEA)
mRRA (ORA)
D
ebGSEA (KPMT)
immune response
leukocyte activation
cell activation
lymphocyte activation
immune system process
B cell receptor signaling pathway
cell periphery
plasma membrane
positive regulation of leukocyte cell-ce
regulation of immune system process
ebGSEA (WT)
protein binding
leukocyte activation
lymphocyte activation
binding
positive regulation of biological proces
membrane
immune system process
regulation of response to stimulus
cell activation
cell periphery
GOmeth
immune response
immune system process
leukocyte activation
cell activation
regulation of immune system process
lymphocyte activation
positive regulation of immune system pro
adaptive immune response
immune effector process
regulation of immune response
HGT
immune system process
response to stimulus
cell periphery
plasma membrane
positive regulation of biological proces
signaling
positive regulation of cellular process
cell communication
immune response
cell activation
mGLM
positive regulation of T cell activation
positive regulation of chronic inflammat
negative regulation of branching involve
positive regulation of synoviocyte proli
positive regulation of superoxide dismut
positive regulation of removal of supero
fractalkine production
regulation of translational initiation b
positive regulation of translational ini
positive regulation of intracellular lip
mRRA (GSEA)
reproduction
negative regulation of transcription by
MAPK cascade
nucleotide binding
nuclear chromosome
chromatin
nuclear chromatin
cell morphogenesis
cell morphogenesis involved in different
transcription regulatory region sequence
mRRA (ORA)
phagocytosis
GTPase binding
regulation of phagocytosis
phosphatidylinositol 3-kinase regulatory
phosphotyrosine residue binding
protein tyrosine kinase binding
receptor tyrosine kinase binding
small GTPase binding
regulation of vesicle-mediated transport
protein phosphorylated amino acid bindin
No. genes 4000 8000 12000 16000
In truth set Both ISP Array Neither
-log10(FDR)

**Supplementary Figure 8. Comparison of gene set testing performance on Gene Ontology (GO) categories for B-cell development data. (A)** Multidimensional scaling plot of 450k array B-cell development data. **(B)** Numbers of differentially methylated CpGs with an adjusted p-value < 0.05, for Stage1 versus Stage 2. The blue bar is the number of significant CpGs that are less methylated in Stage 1 relative to Stage 2, and the red bar is the number that are more methylated; e.g. ~1000 CpGs are less methylated and ~2000 are more methylated in Stage 1, compared to Stage 2. **(C)** Cumulative number of GO terms, as ranked by various methods, that are present in each truth set for Stage 1 versus Stage 2. ISP Terms = immune-system process child terms truth set; Array Terms = top 100 terms from Affymetrix array analysis of the same B-cell development stages. **(D)** Bubble plots of the top 10 GO terms as ranked by various gene set testing methods. The size of the bubble indicates the relative number of genes in the set. The colour of the bubble indicates whether the term is present in either Array (purple) or ISP (green) truth sets, both (red) or neither (blue). ebGSEA (KPMT) = ebGSEA using Known Population Median Test; ebGSEA (WT) = ebGSEA using Wilcoxon Test; GOmeth = GOmeth using either FDR < 0.05 (for contrasts with <5000 significant CpGs) or top 5000 most significant probes; HGT = hypergeometric test; mGLM = methylglm; mRRA (GSEA) = methylRRA using gene set enrichment analysis; mRRA (ORA) = methylRRA using over-representation analysis.

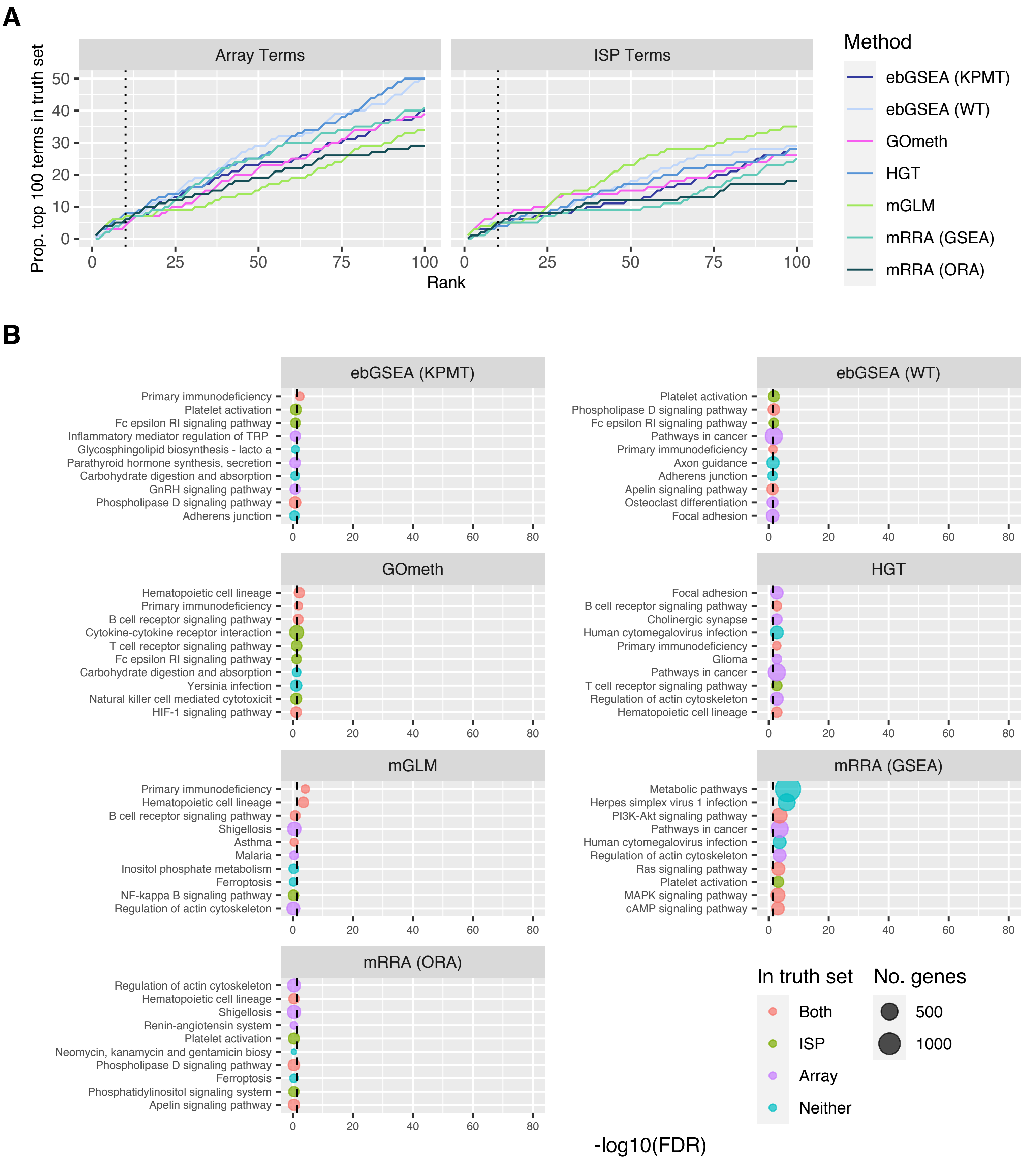
A
Array Terms
ISP Terms
Prop. top 100 terms in truth set
Rank
Method
ebGSEA (KPMT)
ebGSEA (WT)
GOmeth
HGT
mGLM
mRRA (GSEA)
mRRA (ORA)
B
ebGSEA (KPMT)
Primary immunodeficiency
Platelet activation
Fc epsilon RI signaling pathway
Inflammatory mediator regulation of TRP
Glycosphingolipid biosynthesis - lacto a
Parathyroid hormone synthesis, secretion
Carbohydrate digestion and absorption
GnRH signaling pathway
Phospholipase D signaling pathway
Adherens junction
ebGSEA (WT)
Platelet activation
Phospholipase D signaling pathway
Fc epsilon RI signaling pathway
Pathways in cancer
Primary immunodeficiency
Axon guidance
Adherens junction
Apelin signaling pathway
Osteoclast differentiation
Focal adhesion
GOmeth
Hematopoietic cell lineage
Primary immunodeficiency
B cell receptor signaling pathway
Cytokine-cytokine receptor interaction
T cell receptor signaling pathway
Fc epsilon RI signaling pathway
Carbohydrate digestion and absorption
Yersinia infection
Natural killer cell mediated cytotoxicit
HIF-1 signaling pathway
HGT
Focal adhesion
B cell receptor signaling pathway
Cholinergic synapse
Human cytomegalovirus infection
Primary immunodeficiency
Glioma
Pathways in cancer
T cell receptor signaling pathway
Regulation of actin cytoskeleton
Hematopoietic cell lineage
mGLM
Primary immunodeficiency
Hematopoietic cell lineage
B cell receptor signaling pathway
Shigellosis
Asthma
Malaria
Inositol phosphate metabolism
Ferroptosis
NF-kappa B signaling pathway
Regulation of actin cytoskeleton
mRRA (GSEA)
Metabolic pathways
Herpes simplex virus 1 infection
PI3K-Akt signaling pathway
Pathways in cancer
Human cytomegalovirus infection
Regulation of actin cytoskeleton
Ras signaling pathway
Platelet activation
MAPK signaling pathway
cAMP signaling pathway
mRRA (ORA)
Regulation of actin cytoskeleton
Hematopoietic cell lineage
Shigellosis
Renin-angiotensin system
Platelet activation
Neomycin, kanamycin and gentamicin biosy
Phospholipase D signaling pathway
Ferroptosis
Phosphatidylinositol signaling system
Apelin signaling pathway
In truth set
Both
ISP
Array
Neither
No. genes
500
1000
-log10(FDR)

**Supplementary Figure 9. Comparison of gene set testing performance for Stage 1 vs. Stage 2 of B-cell development (KEGG). (A)** Cumulative number of KEGG terms, as ranked by various methods, that are present in each truth set for Stage 1 versus Stage 2. ISP Terms = immune-system process child terms truth set; Array Terms = top 100 terms from Affymetrix array analysis of Stage 1 vs. Stage 2. **(B)** Bubble plots of the top 10 KEGG pathways as ranked by various gene set testing methods. The size of the bubble indicates the relative number of genes in the set. The colour of the bubble indicates whether the term is present in either Array (purple) or ISP (green) truth sets, both (red) or neither (blue). ebGSEA (KPMT) = ebGSEA using Known Population Median Test; ebGSEA (WT) = ebGSEA using Wilcoxon Test; GOmeth = GOmeth using either FDR $< 0.05$ (for contrasts with <5000 significant CpGs) or top 5000 most significant probes; HGT = hypergeometric test; mGLM = methylglm; mRRA (GSEA) = methylRRA using gene set enrichment analysis; mRRA (ORA) = methylRRA using over-representation analysis.

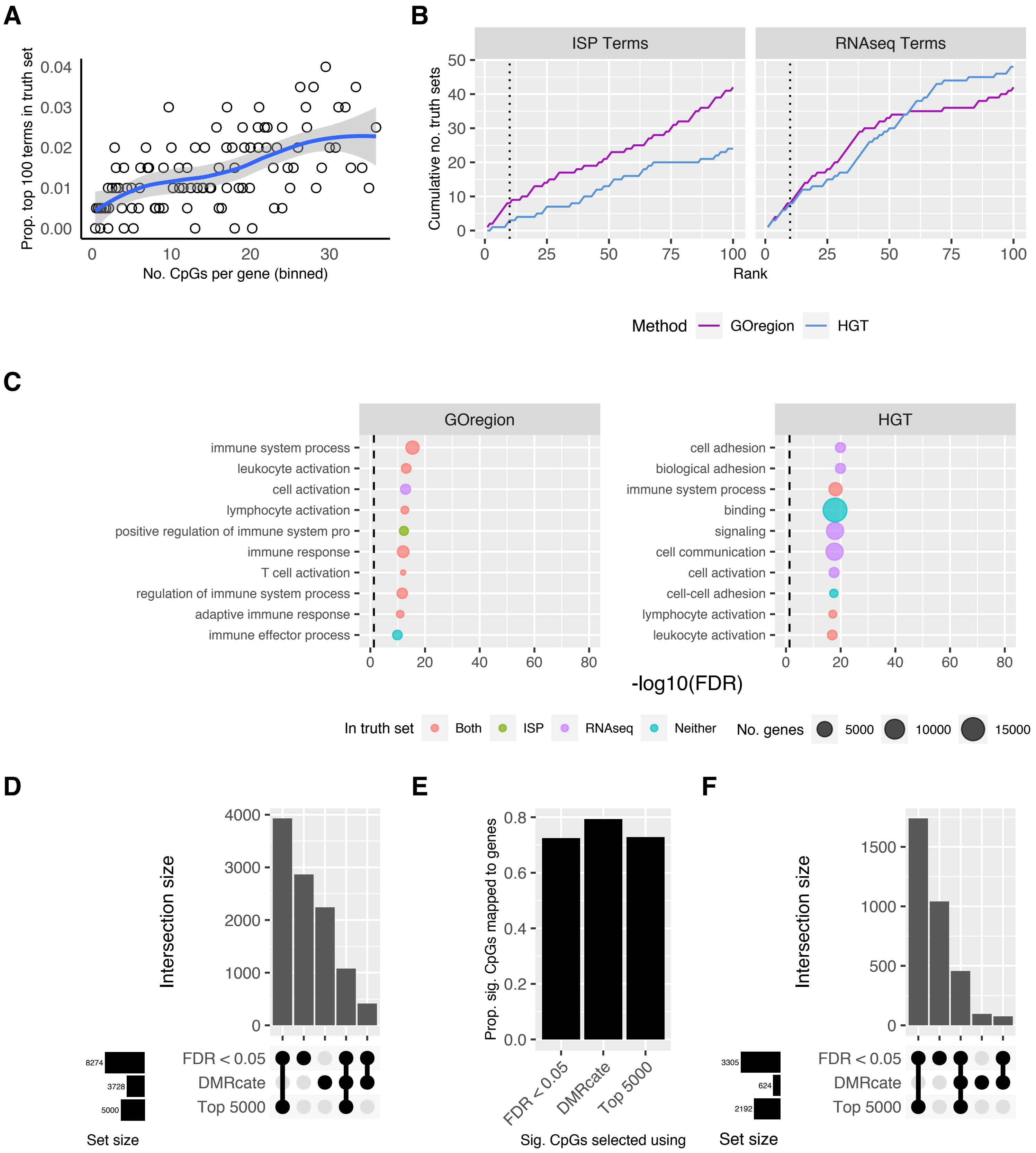
A
Prop. top 100 terms in truth set
No. CpGs per gene (binned)
B
ISP Terms
RNAseq Terms
Cumulative no. truth sets
Rank
Method
GOregion
HGT
C
GOregion
HGT
immune system process
leukocyte activation
cell activation
lymphocyte activation
positive regulation of immune system pro
immune response
T cell activation
regulation of immune system process
adaptive immune response
immune effector process
cell adhesion
biological adhesion
binding
signaling
cell communication
cell-cell adhesion
-log10(FDR)
In truth set
Both
ISP
RNAseq
Neither
No. genes
5000
10000
15000
D
Intersection size
FDR < 0.05
DMRcate
Top 5000
8274
3728
5000
Set size
E
Prop. sig. CpGs mapped to genes
Sig. CpGs selected using
F
3305
624
2192

**Supplementary Figure 10. Evaluation of the performance of GOregion for CD4 T-cells vs. CD8 T-cells. (A)** Bias plot showing that genes that have more CpGs measuring methylation are more likely to have a differentially methylated region. This plot is produced from EPIC array sorted blood cell type data, comparing CD4 T-cells vs. CD8 T-cells. **(B)** Cumulative number of GO terms, as ranked by GOregion and a simple hypergeometric test (HGT), that are present in each truth set for the CD4 T-cells vs. CD8 T-cells comparison. ISP Terms = immune-system process child terms truth set; RNAseq Terms = top 100 terms from RNAseq analysis of the same cell types. **(C)** Bubble plots of the top 10 GO terms as ranked by GOregion and a simple HGT for the CD4 T-cells vs. CD8 T-cells comparison. The size of the bubble indicates the relative number of genes in the set. The colour of the bubble indicates whether the term is present in either RNAseq (purple) or ISP (green) truth sets, both (red) or neither (blue). **(D)** Upset plot showing the characteristics of the CpGs selected as "significant" for the CD4 T-cells vs. CD8 T-cells comparison by a probe-wise differential methylation analysis using a significance cut off (FDR < 0.05), the top 5000 CpGs as ranked by the probe-wise analysis (Top 5000) or the CpGs underlying the filtered *DMRcate* regions (DMRcate). The probe-wise analysis with FDR < 0.05 identified over 8,000 CpGs as "significant" and had the most unique CpGs. Although DMRcate identified the fewest "significant" CpGs (~3,700), over 2,000 were unique to that approach. **(E)** Proportion of "significant" CpGs that are annotated to genes as identified by the three different strategies. **(F)** Upset plot showing the characteristics of the genes that "significant" CpGs are annotated to, as identified by the three different strategies, for the CD4 T-cells vs. CD8 T-cells comparison. CpGs identified by the probe-wise analysis with FDR < 0.05 map to over 3,000 genes. The CpGs identified by DMRcate mapped to ~600 genes, several of which are unique to this approach.

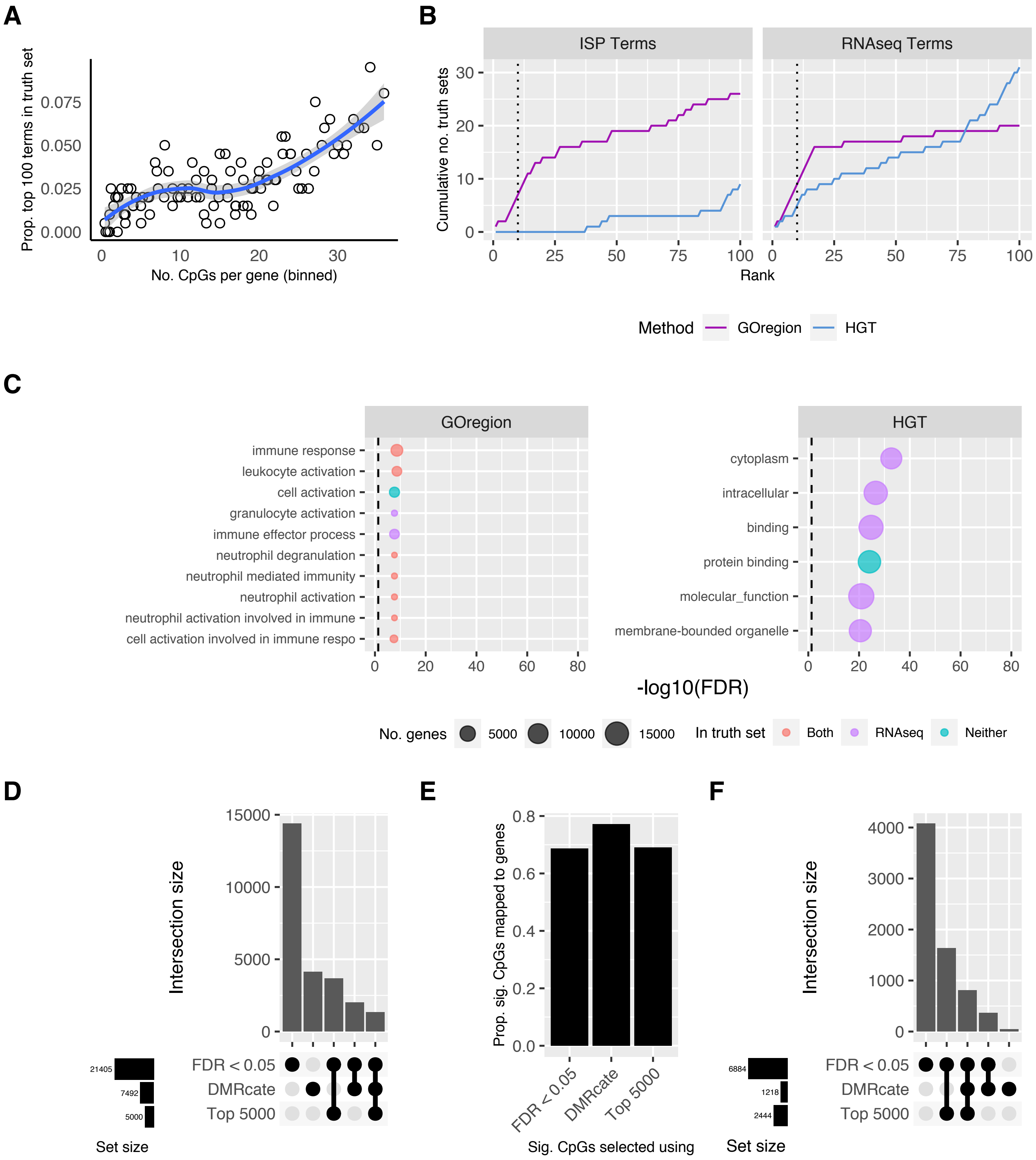
A
Prop. top 100 terms in truth set
0.000
0.025
0.050
0.075
0
10
20
30
No. CpGs per gene (binned)
B
ISP Terms
RNAseq Terms
Cumulative no. truth sets
0
10
20
30
0
25
50
75
100
Rank
Method
GOregion
HGT
C
GOregion
immune response
leukocyte activation
cell activation
granulocyte activation
immune effector process
neutrophil degranulation
neutrophil mediated immunity
neutrophil activation
neutrophil activation involved in immune
cell activation involved in immune respo
HGT
cytoplasm
intracellular
binding
protein binding
molecular_function
membrane-bounded organelle
0
20
40
60
80
-log10(FDR)
No. genes
5000
10000
15000
In truth set
Both
RNAseq
Neither
D
Intersection size
15000
10000
5000
0
21405
7492
5000
Set size
FDR < 0.05
DMRcate
Top 5000
E
Prop. sig. CpGs mapped to genes
0.0
0.2
0.4
0.6
0.8
FDR < 0.05
DMRcate
Top 5000
Sig. CpGs selected using
F
Intersection size
4000
3000
2000
1000
0
6884
1218
2444
Set size
FDR < 0.05
DMRcate
Top 5000

**Supplementary Figure 11. Evaluation of the performance of GOregion for monocytes vs. neutrophils. (A)** Bias plot showing that genes that have more CpGs measuring methylation are more likely to have a differentially methylated region. This plot is produced from EPIC array sorted blood cell type data, comparing monocytes vs. neutrophils. **(B)** Cumulative number of GO terms, as ranked by GOregion and a simple hypergeometric test (HGT), that are present in each truth set for the monocytes vs. neutrophils comparison. ISP Terms = immune-system process child terms truth set; RNAseq Terms = top 100 terms from RNAseq analysis of the same cell types. **(C)** Bubble plots of the top 10 GO terms as ranked by GOregion and a simple HGT for the monocytes vs. neutrophils comparison. The size of the bubble indicates the relative number of genes in the set. The colour of the bubble indicates whether the term is present in either RNAseq (purple) or ISP (green) truth sets, both (red) or neither (blue). **(D)** Upset plot showing the characteristics of the CpGs selected as "significant" for the monocytes vs. neutrophils comparison by a probe-wise differential methylation analysis using a significance cut off (FDR < 0.05), the top 5000 CpGs as ranked by the probe-wise analysis (Top 5000) or the CpGs underlying the filtered *DMRcate* regions (DMRcate). The probe-wise analysis with FDR < 0.05 identified over 21,000 CpGs as "significant" and had the most unique CpGs. Although DMRcate identified fewer CpGs (~7500), ~3500 of them were unique to that approach. **(E)** Proportion of "significant" CpGs that are annotated to genes as identified by the three different strategies. **(F)** Upset plot showing the characteristics of the genes that "significant" CpGs are annotated to, as identified by the three different strategies, for the monocytes vs. neutrophils comparison. CpGs identified by the probe-wise analysis with FDR < 0.05 map to over 6,000 genes. The CpGs identified by DMRcate mapped to ~1200, several of which are unique to this approach.

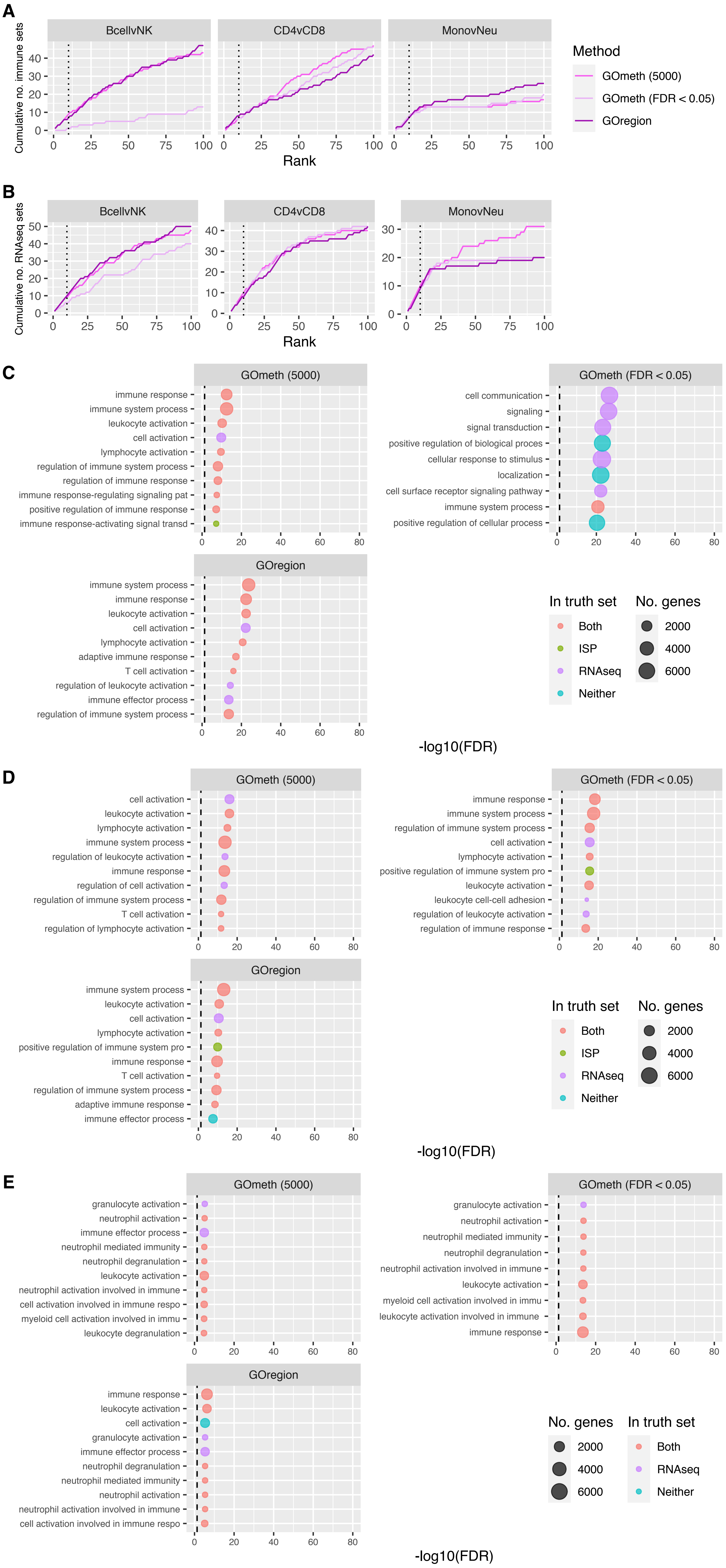
A
BcellvNK
CD4vCD8
MonovNeu
Cumulative no. immune sets
Rank
Method
GOmeth (5000)
GOmeth (FDR < 0.05)
GOregion
B
BcellvNK
CD4vCD8
MonovNeu
Cumulative no. RNAseq sets
Rank
C
GOmeth (5000)
immune response
immune system process
leukocyte activation
cell activation
lymphocyte activation
regulation of immune system process
regulation of immune response
immune response-regulating signaling pat
positive regulation of immune response
immune response-activating signal transd
GOmeth (FDR < 0.05)
cell communication
signaling
signal transduction
positive regulation of biological proces
cellular response to stimulus
localization
cell surface receptor signaling pathway
immune system process
positive regulation of cellular process
GOregion
immune system process
immune response
leukocyte activation
cell activation
lymphocyte activation
adaptive immune response
T cell activation
regulation of leukocyte activation
immune effector process
regulation of immune system process
In truth set
Both
ISP
RNAseq
Neither
No. genes
2000
4000
6000
-log10(FDR)
D
GOmeth (5000)
cell activation
leukocyte activation
lymphocyte activation
immune system process
regulation of leukocyte activation
immune response
regulation of cell activation
regulation of immune system process
T cell activation
regulation of lymphocyte activation
GOmeth (FDR < 0.05)
immune response
immune system process
regulation of immune system process
cell activation
lymphocyte activation
positive regulation of immune system pro
leukocyte activation
leukocyte cell-cell adhesion
regulation of leukocyte activation
regulation of immune response
GOregion
immune system process
leukocyte activation
cell activation
lymphocyte activation
positive regulation of immune system pro
immune response
T cell activation
regulation of immune system process
adaptive immune response
immune effector process
In truth set
Both
ISP
RNAseq
Neither
No. genes
2000
4000
6000
-log10(FDR)
E
GOmeth (5000)
granulocyte activation
neutrophil activation
immune effector process
neutrophil mediated immunity
neutrophil degranulation
leukocyte activation
neutrophil activation involved in immune
cell activation involved in immune respo
myeloid cell activation involved in immu
leukocyte degranulation
GOmeth (FDR < 0.05)
granulocyte activation
neutrophil activation
neutrophil mediated immunity
neutrophil degranulation
neutrophil activation involved in immune
leukocyte activation
myeloid cell activation involved in immu
leukocyte activation involved in immune
immune response
GOregion
immune response
leukocyte activation
cell activation
granulocyte activation
immune effector process
neutrophil degranulation
neutrophil mediated immunity
neutrophil activation
neutrophil activation involved in immune
cell activation involved in immune respo
No. genes
2000
4000
6000
In truth set
Both
RNAseq
Neither
-log10(FDR)

**Supplementary Figure 12. Comparison of probe-wise and region-wise analyses in the gene set testing context for sorted blood cell types data. (A)** Cumulative number of GO terms, as ranked by various analysis strategies, that are present in the immune truth set for all comparisons. ISP Terms = immune-system process child terms truth set. **(B)** Cumulative number of GO terms, as ranked by various methods, that are present in the RNAseq truth set for all comparisons. RNAseq Terms = top 100 terms from RNAseq analysis of the same cell types. **(C)** Bubble plots of the top 10 GO terms as ranked by various analysis strategies for the B-cell vs. NK comparison. The size of the bubble indicates the relative number of genes in the set. The colour of the bubble indicates whether the term is present in either RNAseq (purple) or ISP (green) truth sets, both (red) or neither (blue). **(D)** Bubble plots of the top 10 GO terms as ranked by various analysis strategies for the CD4 T-cells vs. CD8 T-cells comparison. **(E)** Bubble plots of the top 10 GO terms as ranked by various analysis strategies for the monocytes vs. neutrophils comparison. GOmeth (5000) = GOmeth using top 5000 most significant probes; GOmeth (FDR < 0.05) = GOmeth using all probes significant at FDR < 0.05; GOregion = GOregion using all selected region coordinates.

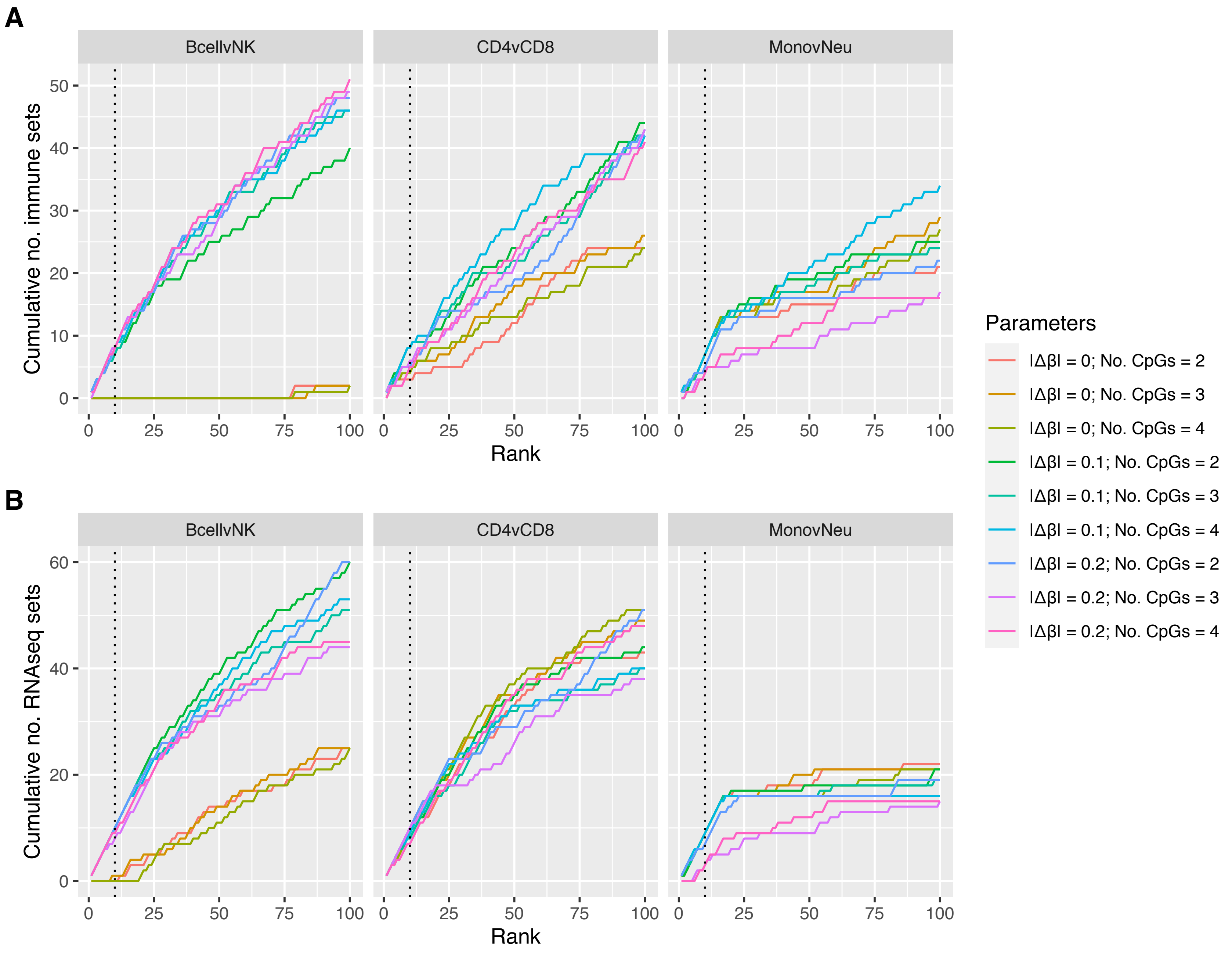

A
BcellvNK
CD4vCD8
MonovNeu
Cumulative no. immune sets
Rank
B
Cumulative no. RNAseq sets
Parameters
|Δβ| = 0; No. CpGs = 2
|Δβ| = 0; No. CpGs = 3
|Δβ| = 0; No. CpGs = 4
|Δβ| = 0.1; No. CpGs = 2
|Δβ| = 0.1; No. CpGs = 3
|Δβ| = 0.1; No. CpGs = 4
|Δβ| = 0.2; No. CpGs = 2
|Δβ| = 0.2; No. CpGs = 3
|Δβ| = 0.2; No. CpGs = 4

**Supplementary Figure 13. Effect of DMR filtering on gene set testing for sorted blood cell types data. (A)** Cumulative number of GO terms, as ranked by GOregion analysis of DMRs filtered using various parameters, that are present in the immune truth set for all comparisons. ISP Terms = immune-system process child terms truth set. **(B)** Cumulative number of GO terms, as ranked by GOregion analysis of DMRs filtered using various parameters, that are present in the RNAseq truth set for all comparisons. RNAseq Terms = top 100 terms from RNAseq analysis of the same cell types. $|\Delta\beta|$ = mean methylation difference across region; No. CpGs = number of CpGs underlying region.

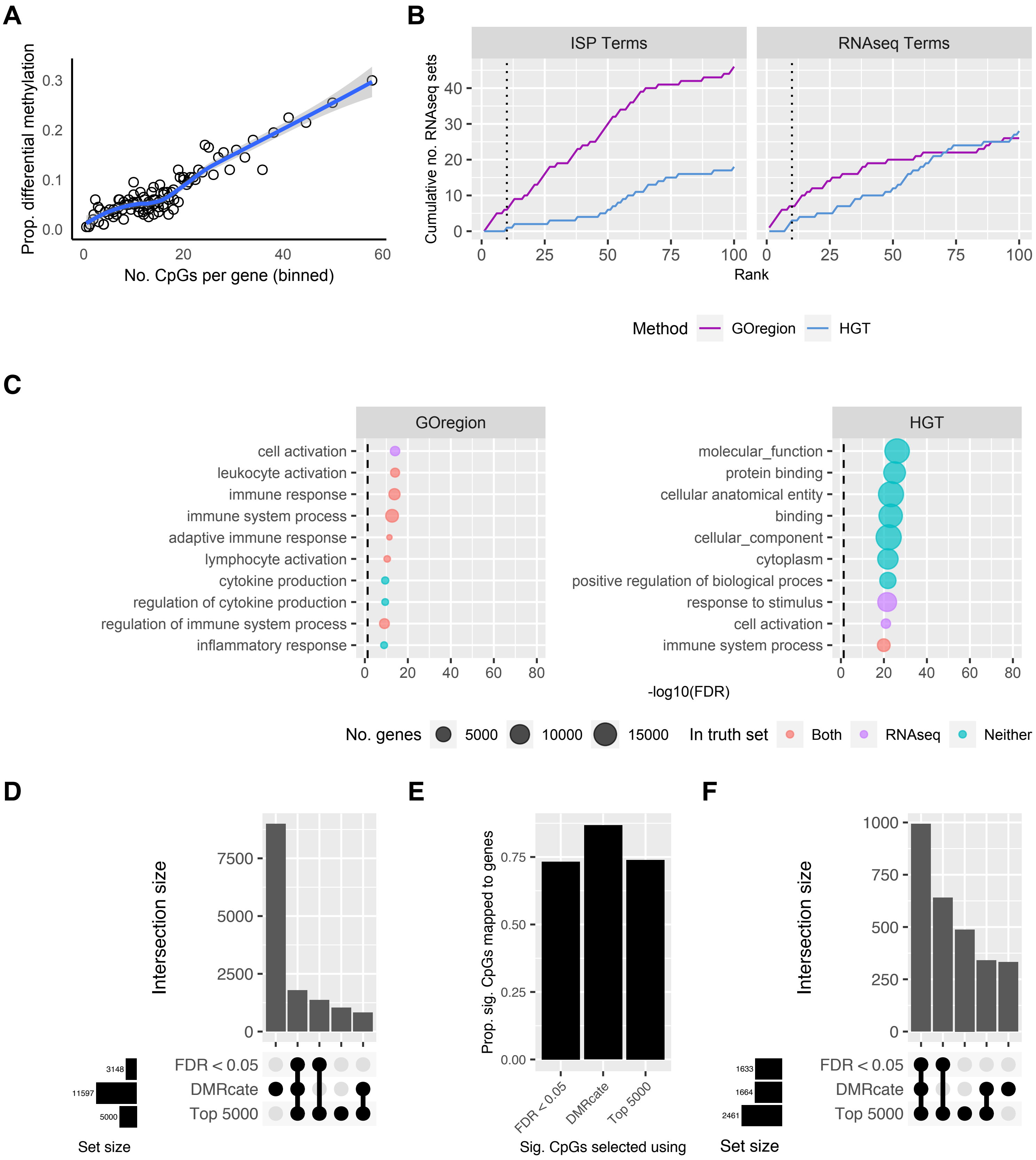
A
Prop. differential methylation
No. CpGs per gene (binned)
B
ISP Terms
RNAseq Terms
Cumulative no. RNAseq sets
Rank
Method
GOregion
HGT
C
GOregion
cell activation
leukocyte activation
immune response
immune system process
adaptive immune response
lymphocyte activation
cytokine production
regulation of cytokine production
regulation of immune system process
inflammatory response
HGT
molecular_function
protein binding
cellular anatomical entity
binding
cellular_component
cytoplasm
positive regulation of biological proces
response to stimulus
cell activation
immune system process
-log10(FDR)
No. genes
5000
10000
15000
In truth set
Both
RNAseq
Neither
D
Intersection size
FDR < 0.05
DMRcate
Top 5000
3148
11597
5000
Set size
E
Prop. sig. CpGs mapped to genes
FDR < 0.05
DMRcate
Top 5000
Sig. CpGs selected using
F
Intersection size
FDR < 0.05
DMRcate
Top 5000
1633
1664
2461
Set size

**Supplementary Figure 14. Evaluation of the performance of GOregion on B-cell development data. (A)** Bias plot showing that genes with more measured CpGs are more likely to have a differentially methylated region (DMR). This plot is produced from 450K array B-cell development data, comparing Stage 1 vs. Stage 2. **(B)** Cumulative number of GO terms, as ranked by GOregion and a simple hypergeometric test (HGT), that are present in each truth set for the Stage 1 versus Stage 2 comparison. ISP Terms = immune-system process child terms truth set; Array Terms = top 100 terms from Affymetrix array analysis of the same B-cell development stages. **(C)** Bubble plots of the top 10 GO terms as ranked by GOregion and a simple HGT for the Stage 1 versus Stage 2 comparison. The size of the bubble indicates the relative number of genes in the set. The colour of the bubble indicates whether the term is present in either Array (purple) or ISP (green) truth sets, both (red) or neither (blue). **(D)** Upset plot showing the characteristics of the CpGs selected as "significant" for the Stage 1 versus Stage 2 comparison by a probe-wise differential methylation analysis using a significance cut off (FDR < 0.05), the top 5000 CpGs as ranked by the probe-wise analysis (Top 5000) or the CpGs underlying the filtered *DMRcate* regions (DMRcate). The *DMRcate* analysis identified over 11,000 CpGs as "significant" and had the most unique CpGs (>7500). **(E)** Proportion of "significant" CpGs that are annotated to genes as identified by the three different strategies. **(F)** Upset plot showing the characteristics of the genes that "significant" CpGs are annotated to, as identified by the three different strategies, for the Stage 1 versus Stage 2 comparison.

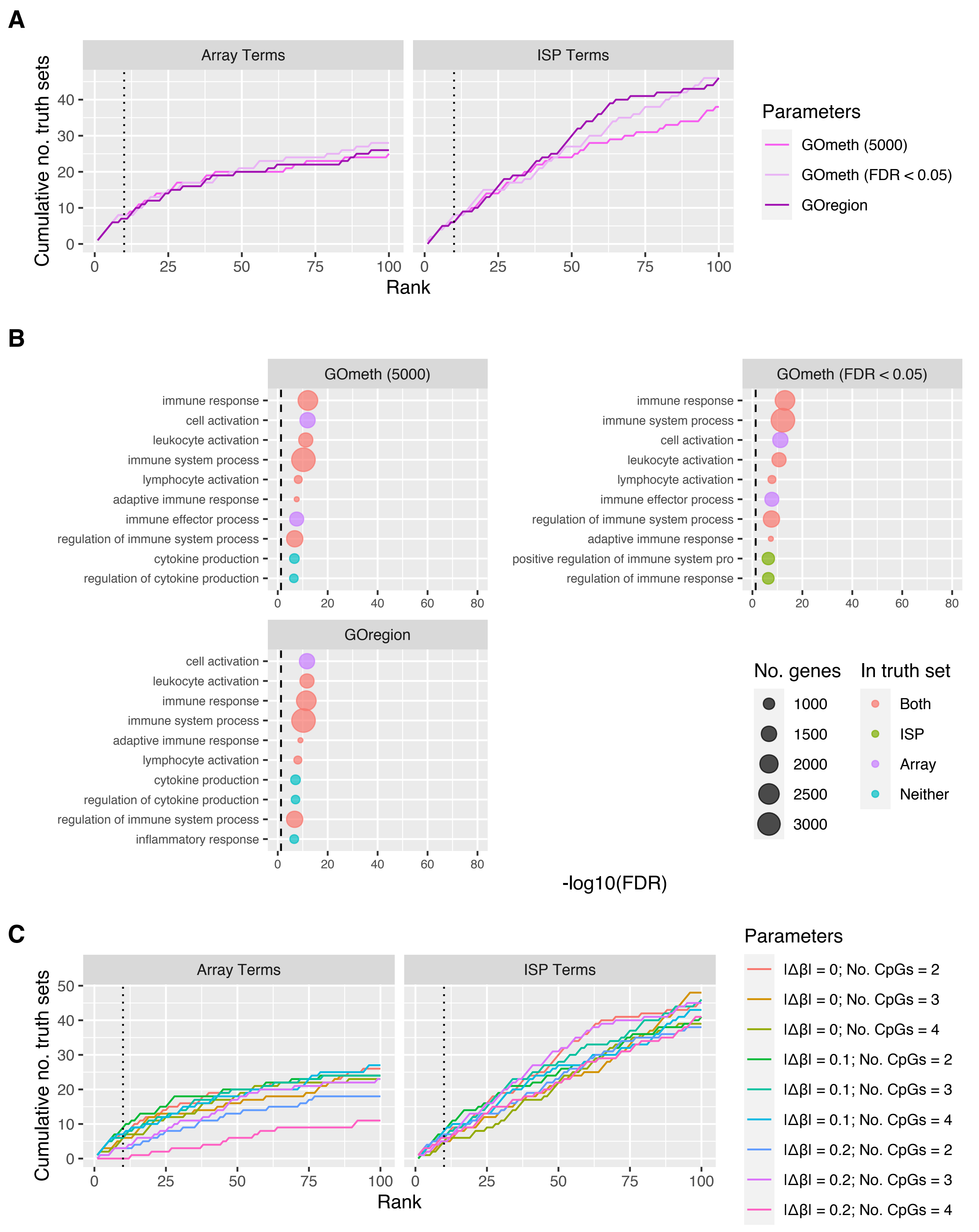

A
Array Terms
ISP Terms
Cumulative no. truth sets
Rank
Parameters
GOmeth (5000)
GOmeth (FDR < 0.05)
GOregion
B
GOmeth (5000)
immune response
cell activation
leukocyte activation
immune system process
lymphocyte activation
adaptive immune response
immune effector process
regulation of immune system process
cytokine production
regulation of cytokine production
GOmeth (FDR < 0.05)
immune response
immune system process
cell activation
leukocyte activation
lymphocyte activation
immune effector process
regulation of immune system process
adaptive immune response
positive regulation of immune system pro
regulation of immune response
GOregion
cell activation
leukocyte activation
immune response
immune system process
adaptive immune response
lymphocyte activation
cytokine production
regulation of cytokine production
regulation of immune system process
inflammatory response
-log10(FDR)
No. genes
1000
1500
2000
2500
3000
In truth set
Both
ISP
Array
Neither
C
Array Terms
ISP Terms
Cumulative no. truth sets
Rank
Parameters
|Δβ| = 0; No. CpGs = 2
|Δβ| = 0; No. CpGs = 3
|Δβ| = 0; No. CpGs = 4
|Δβ| = 0.1; No. CpGs = 2
|Δβ| = 0.1; No. CpGs = 3
|Δβ| = 0.1; No. CpGs = 4
|Δβ| = 0.2; No. CpGs = 2
|Δβ| = 0.2; No. CpGs = 3
|Δβ| = 0.2; No. CpGs = 4

**Supplementary Figure 15. Comparison of probe-wise and region-wise analyses in the gene set testing context using B-cell development data. (A)** Cumulative number of GO terms, as ranked by various analysis strategies, that are present in each truth set for Stage 1 vs. Stage 2. ISP Terms = immune-system process child terms truth set; Array Terms = top 100 terms from Affymetrix array analysis of Stage 1 vs. Stage 2. GOmeth (5000) = GOmeth using top 5000 most significant probes; GOmeth (FDR $< 0.05$) = GOmeth using all probes significant at FDR $< 0.05$; GOregion = GOregion using all selected region coordinates. **(B)** Bubble plots of the top 10 GO terms as ranked by various analysis strategies for the Stage 1 vs. Stage 2 comparison. The size of the bubble indicates the relative number of genes in the set. The colour of the bubble indicates whether the term is present in either Array (purple) or ISP (green) truth sets, both (red) or neither (blue). **(C)** Effect of DMR filtering on gene set testing. Cumulative number of GO terms, as ranked by GOregion analysis of DMRs filtered using various parameters, that are present in each truth set for the Stage 1 vs. Stage 2 comparison. ISP Terms = immune-system process child terms truth set. Array Terms = top 100 terms from Affymetrix array analysis of the same B-cell development stages. $|\Delta\beta|$ = mean methylation difference across region; No. CpGs = number of CpGs underlying region.
